# Persisting perceptual abnormalities in psychedelic users are associated with conditioned hallucinations and impaired sensory processing

**DOI:** 10.64898/2026.08.10.743999

**Authors:** Maximillian S. Greenwald, Peter T. Waade, Eren Kafadar, K. Alexandria Bond, Dominik Firisz, Samuel W. Nehrer, Saida Ibragimova, Albert R. Powers

**Author notes:** Correspondence should be addressed to: Albert R. Powers, M.D., Ph.D., The Connecticut Mental Health Center, Rm. S109 34 Park Street, New Haven, CT 06519.

## Abstract

Serotonergic psychedelics (SP) are increasingly used in clinical research and naturalistic settings, but their psychotic-like side effects, including persisting perceptual abnormalities (PPAs), are poorly understood. Psychosis-associated hallucinations are associated with susceptibility to conditioned hallucinations and computationally-estimated overweighting of perceptual expectations, or priors. However, SPs are widely argued to reduce prior weighting. We surveyed 186 naturalistic SP users on prior SP use, SP-associated PPA history, and current PPAs. Participants completed the visual conditioned hallucinations (VCH) task, in which conditioning induces perception of absent stimuli. Behavioral data were used to fit parameters of a computational model to estimate latent states driving percepts and responses. Past and current PPAs were associated with younger age at first use and higher SP doses, lower visual thresholds, higher VCH rate and confidence, and reduced sensory discrimination. Among model parameters, however, only reduced decision precision tracked both measures and mediated the dose-PPA relationship; relative prior weighting rose equivocally, as expected when priors and sensory evidence gain precision together. SP-related PPAs may therefore arise from a noisy visual system biased toward detection, in which priors act as templates that convert sensory noise into expected percepts. These findings may point to a tractable model for how psychotic-like perception emerges.

## Introduction

A growing body of research suggests that treatment with serotonergic psychedelics (SPs) may reduce symptoms across many psychiatric disorders,^1^ with effects that persist months to years^2^ after dosing. Cross-sectional and prospective online studies suggest that even extra-clinical, naturalistic SP use is associated with persisting protective effects.^3,4^ However, there is also evidence that SPs induce acute^5^ and persisting^3,6–13^ psychotic-like experiences. These psychotic-like experiences differ from psychotic symptoms in their subtle nature and distinct phenomenology, characterized by experiences like easy absorption in one’s imagination^12^ and minor visual imagery or phantom-like persisting perceptual abnormalities (PPAs).^6^ These experiences also resemble visual distortions frequently reported in clinical high risk for psychosis (CHR-P),^14^ suggesting they may meaningfully relate to psychosis risk. How these effects relate to the mechanisms driving primary psychotic illness remains poorly understood. Studying the PPAs associated with SP use may serve as a way of delineating the pathways linking SP exposure to the emergence of psychotic symptoms like hallucinations and the persistence of both therapeutic and psychotogenic effects.

PPAs following SP use have long been studied within the context of Hallucinogen Persisting Perception Disorder (HPPD), a DSM-V diagnosis requiring recurrence of SP-induced percepts causing clinically significant impairment and distress.^15^ Using these criteria, HPPD is rare (1-5% of SP users). However, expanding PPA assessment to include subtle and non-distressing/impairing effects has demonstrated that PPAs may be the single most common persisting side effect of SP use. Between ∼25%^10^ and ∼50%,^6,7^ of SP users endorse current PPAs, and prospective cohort studies have found between ∼33%^3,9^ and ∼75%^3^ of users endorsing PPAs following a single SP use. Effects are most typically reported days^16^ to weeks^3,9^ later, with evidence suggesting that PPAs fade with time (e.g. 25% down to 3% in one year).^10^ Objective, neurally-linked measures associated with PPAs to date have also focused exclusively on individuals diagnosed with HPPD. Findings associated with HPPD include reduced occipital coherence with distant brain regions, fast alpha rhythm and shortened visual-evoked potential latency, increased occipital delta power, and symptom exacerbation on treatment with risperidone (a 5-HT2AR antagonist).^15^ A consistent explanation offered for these findings is disinhibition/hyperactivation in the visual pathway.^17^

The acute perceptual abnormalities and therapeutic effects arising from SP use have been explained within a hierarchical predictive processing theoretical framework. In this framework, percepts result from a combination of incoming, bottom-up sensory information with top-down prior expectations (i.e., *priors*) weighted by the relative reliability—or precision—of the information.^18^ The most popular computational theory leveraging this framework to study SP action (Relaxed Beliefs Under pSychedelics; REBUS) argues that SPs reduce prior precision and allow aberrant sensory noise to dominate perception.^18^ REBUS offers an intuitive explanation for SPs’ therapeutic efficacy and hallucinatory phenomena, which consist of elementary visual features.^19^ It is further supported by EEG^20,21^ and fMRI^22,23^ studies that report reduced information flow from hierarchically higher to lower brain regions using various modeling approaches (but see two contrary high-quality fMRI studies).^24,25^

Yet, if SPs’ *acute* perceptual abnormalities occur via reduced prior precision, how their *persisting* perceptual abnormalities might occur remains a key open question. Do persisting perceptual abnormalities result from persistent reductions in prior weighting and propagation of bottom-up noise, or might they reflect compensatory *increases* in prior weighting? Supporting this latter account, computational modeling of behavior^26–29^ and EEG^20,30^ consistently suggest that hallucinations result from *overly* precise priors.^31^ This is directly demonstrated by consistent associations between hallucinatory symptoms and susceptibility to conditioned hallucinations.^26–29^ Recent work has argued that this prior overweighting in psychosis-related hallucinations may arise as a compensatory response to sensory noise arising from acute SP action.^31^ However, no work has yet directly tested whether PPAs related to SP use are due to such a compensatory process or to ongoing propagation of unconstrained bottom-up noise.

Here, we explore the computational and behavioral signatures of perceptual inference in SP-related PPAs.^26–29^ We recruited an online sample of SP users and non-users who answered detailed questions regarding their SP use, history of SP-associated PPAs, and current PPAs before then completing the Visual Conditioned Hallucinations (VCH) task—a visual variant on a behavioral task sensitive to hallucinatory processes in the psychosis spectrum. We examined how orthogonal measures of SP exposure vary with current PPAs and risk of lifetime SP-associated PPAs as well as how both vary with *in-vivo* hallucinatory behavior in the VCH and model-derived estimates of individuals’ perceptual inference, including weighting of priors relative to sensory evidence. We predicted that SP-related PPAs would be associated with fewer conditioned hallucinations and a lower prior:sensory evidence precision ratio if they were driven by persisting prior *hypo*-precision as described in the REBUS theory and the opposite if driven by compensatory prior *hyper*-precision.

## Methods

### Procedures

All procedures were approved by the Yale University Institutional Review Board (IRB) / Human Research Protection Program (HRPP) and were conducted in accordance with the Declaration of Helsinki. Participants signed an E-consent form via a public survey link after correctly passing a CAPTCHA test through Yale’s instantiation of the web-based and HIPAA-secure Research Electronic Data Capture platform (REDCap@Yale).^32^ They were then directed to a screening form to ensure all participants met inclusion criteria and to rule out automated responses, repeat participation, and fraud (see **Supplementary Materials**). All included participants then completed the primary study questionnaires and VCH task within 72 hours. Those who failed to finish within 72 hours were notified and allowed to restart.

All participants provided informed consent and received monetary compensation if they completed all study procedures and passed quality checks described above.

### Participants

Individuals endorsing prior use of supra-threshold doses of any SP (compounds that primarily agonize the Serotonin 2A receptor)^33^ as well as non-users (for sensitivity analyses) were recruited between August of 2024 through January of 2026 from several sources: SP online community discussion forums including Bluelight.org, Shroomery.org, DMT Nexus, r/Drugs, r/Psychonaut, r/Shrooms, r/Mescaline, r/Ayahuasca, r/DMT, r/5MeODMT, r/HPPD, and r/LSD; as well as snowball recruitment primarily from prior participants^26^ and university students (**Table 1**).

**Table 1.** Demographics, IQ proxies, psychiatric history, and substance use. Sample compositions are compared separately between SP users endorsing and denying a history of SP-associated PPAs using a modified 12-item questionnaire (middle table; n = 186) and those endorsing vs. denying current PPAs (right table; n = 130). P values indicate the result of a Mann-Whitney U test in the case of continuous variables and a 2×2 Chi-square test of independence (comparing that level of the variable between groups) in the case of categorical variables with more than five observations at both levels and a Fisher’s Exact test otherwise.

| Characteristic | PPA (-) | PPA (+) | Total | P-value | CAPS(-) | CAPS(+) | Total | P-value |
| --- | --- | --- | --- | --- | --- | --- | --- | --- |
| **Psychiatric History** |  |  |  |  |  |  |  |  |
| Any Diagnosis | 23<br>(41.8%) | 46 (35.1%) | 69<br>(37.1%) | 0.486 | 24 (38.7%) | 26 (38.2%) | 50<br>(38.5%) | 1.000 |
| Psychotic Spectrum | 3 (5.5%) | 4 (3.1%) | 7 (3.8%) | 0.424 | 0 (0.0%) | 2 (2.9%) | 2 (1.5%) | 0.497 |
| Schizophrenia Spectrum | 3 (5.5%) | 4 (3.1%) | 7 (3.8%) | 0.424 | 0 (0.0%) | 2 (2.9%) | 2 (1.5%) | 0.497 |
| Unipolar Mood Disorder | 11<br>(20.0%) | 19 (14.5%) | 30<br>(16.1%) | 0.477 | 12 (19.4%) | 11 (16.2%) | 23<br>(17.7%) | 0.807 |
| Bipolar Disorder (no psychosis) | 1 (1.8%) | 4 (3.1%) | 5 (2.7%) | 1.000 | 1 (1.6%) | 3 (4.4%) | 4 (3.1%) | 0.621 |
| Substance Use Disorder | 1 (1.8%) | 8 (6.1%) | 9 (4.8%) | 0.285 | 1 (1.6%) | 5 (7.4%) | 6 (4.6%) | 0.211 |
| Anxiety Disorder | 10<br>(18.2%) | 22 (16.8%) | 32<br>(17.2%) | 0.987 | 12 (19.4%) | 11 (16.2%) | 23<br>(17.7%) | 0.807 |
| Obsessive-Compulsive Disorder (OCD) | 1 (1.8%) | 1 (0.8%) | 2 (1.1%) | 0.505 | 1 (1.6%) | 1 (1.5%) | 2 (1.5%) | 1.000 |
| Trauma-Related Disorder (PTSD/c-PTSD) | 7 (12.7%) | 9 (6.9%) | 16 (8.6%) | 0.251 | 5 (8.1%) | 6 (8.8%) | 11<br>(8.5%) | 1.000 |
| Eating Disorder | 0 (0.0%) | 1 (0.8%) | 1 (0.5%) | 1.000 | 0 (0.0%) | 1 (1.5%) | 1 (0.8%) | 1.000 |
| Personality Disorder | 1 (1.8%) | 4 (3.1%) | 5 (2.7%) | 1.000 | 2 (3.2%) | 1 (1.5%) | 3 (2.3%) | 0.605 |
| Autism Spectrum Disorder | 1 (1.8%) | 6 (4.6%) | 7 (3.8%) | 0.676 | 3 (4.8%) | 4 (5.9%) | 7 (5.4%) | 1.000 |
| Attention Deficit Hyperactivity Disorder | 10<br>(18.2%) | 17 (13.0%) | 27<br>(14.5%) | 0.489 | 11 (17.7%) | 9 (13.2%) | 20<br>(15.4%) | 0.640 |
| Sleep Disorder | 2 (3.6%) | 5 (3.8%) | 7 (3.8%) | 1.000 | 3 (4.8%) | 2 (2.9%) | 5 (3.8%) | 0.669 |
| **Psychiatric Medications** |  |  |  |  |  |  |  |  |
| Any Medication | 17<br>(30.9%) | 23 (17.6%) | 40<br>(21.5%) | 0.068 | 17 (27.4%) | 11 (16.2%) | 28<br>(21.5%) | 0.179 |
| Antipsychotic | 1 (1.8%) | 2 (1.5%) | 3 (1.6%) | 1.000 | 2 (3.2%) | 0 (0.0%) | 2 (1.5%) | 0.226 |
| Antidepressant | 6 (10.9%) | 10 (7.6%) | 16 (8.6%) | 0.567 | 6 (9.7%) | 6 (8.8%) | 12<br>(9.2%) | 1.000 |
| Stimulant | 8 (14.5%) | 8 (6.1%) | 16 (8.6%) | 0.084 | 6 (9.7%) | 2 (2.9%) | 8 (6.2%) | 0.150 |
| Benzodiazepine | 1 (1.8%) | 2 (1.5%) | 3 (1.6%) | 1.000 | 2 (3.2%) | 1 (1.5%) | 3 (2.3%) | 0.605 |
| Anxiolytic (Non-Benzodiazepine) | 3 (5.5%) | 1 (0.8%) | 4 (2.2%) | 0.078 | 3 (4.8%) | 1 (1.5%) | 4 (3.1%) | 0.347 |
| Sedative (Non-Benzodiazepine) | 3 (5.5%) | 2 (1.5%) | 5 (2.7%) | 0.155 | 3 (4.8%) | 0 (0.0%) | 3 (2.3%) | 0.106 |
| Opioid Antagonist | 0 (0.0%) | 2 (1.5%) | 2 (1.1%) | 1.000 | 0 (0.0%) | 2 (2.9%) | 2 (1.5%) | 0.497 |
| **Other Substance Use (Past Month)** |  |  |  |  |  |  |  |  |
| Any Substance Use (Past Month) | 32<br>(58.2%) | 93 (71.0%) | 125<br>(67.2%) | 0.127 | 43 (69.4%) | 43 (63.2%) | 86<br>(66.2%) | 0.582 |
| Alcohol Use (Past Month) | 26<br>(47.3%) | 76 (58.0%) | 102<br>(54.8%) | 0.237 | 33 (53.2%) | 38 (55.9%) | 71<br>(54.6%) | 0.899 |
| Sedative-Hynotic Use (Past Month) | 3 (5.5%) | 14 (10.7%) | 17 (9.1%) | 0.395 | 5 (8.1%) | 2 (2.9%) | 7 (5.4%) | 0.257 |
| Opioid Use (Past Month) | 1 (1.8%) | 6 (4.6%) | 7 (3.8%) | 0.676 | 2 (3.2%) | 1 (1.5%) | 3 (2.3%) | 0.605 |
| Cannabis Use (Past Month) | 15<br>(27.3%) | 34 (26.0%) | 49<br>(26.3%) | 0.997 | 17 (27.4%) | 14 (20.6%) | 31<br>(23.8%) | 0.480 |
| Atypical Psychedelics Use<br>(Past Month) | 4 (7.3%) | 13 (9.9%) | 17 (9.1%) | 0.769 | 3 (4.8%) | 5 (7.4%) | 8 (6.2%) | 0.720 |
| Stimulants Use (Past Month) | 2 (3.6%) | 16 (12.2%) | 18 (9.7%) | 0.125 | 2 (3.2%) | 4 (5.9%) | 6 (4.6%) | 0.682 |
| **Other Substance Use History (Lifetime)** |  |  |  |  |  |  |  |  |
| Any Substance Use (Lifetime) | 55<br>(100.0%) | 131 (100.0%) | 186<br>(100.0%) | 1.000 | 62<br>(100.0%) | 68<br>(100.0%) | 130<br>(100.0%) | 1.000 |
| Alcohol Use (Lifetime) | 50<br>(90.9%) | 116 (88.5%) | 166<br>(89.2%) | 0.830 | 55 (88.7%) | 58 (85.3%) | 113<br>(86.9%) | 0.752 |
| Sedative-Hypnotic Use<br>(Lifetime) | 16<br>(29.1%) | 57 (43.5%) | 73<br>(39.2%) | 0.094 | 22 (35.5%) | 24 (35.3%) | 46<br>(35.4%) | 1.000 |
| Opioid Use (Lifetime) | 12<br>(21.8%) | 40 (30.5%) | 52<br>(28.0%) | 0.303 | 16 (25.8%) | 15 (22.1%) | 31<br>(23.8%) | 0.768 |
| Cannabis Use (Lifetime) | 19<br>(34.5%) | 43 (32.8%) | 62<br>(33.3%) | 0.955 | 24 (38.7%) | 20 (29.4%) | 44<br>(33.8%) | 0.351 |
| Atypical Psychedelics Use<br>(Lifetime) | 36<br>(65.5%) | 82 (62.6%) | 118<br>(63.4%) | 0.839 | 36 (58.1%) | 44 (64.7%) | 80<br>(61.5%) | 0.551 |
| Stimulants Use (Lifetime) | 24<br>(43.6%) | 74 (56.5%) | 98<br>(52.7%) | 0.150 | 26 (41.9%) | 33 (48.5%) | 59<br>(45.4%) | 0.563 |

Participants were allowed to complete the study only if they met the following inclusion criteria: 18-65 years old, fluent in English, access to a laptop or computer with stable internet and headphones, no history of seizures or neurological disorders causing cognitive impairment, and correct responses to 3/9 Raven’s progressive matrices.^34^ SP users were excluded if they had used non-5-HT2AR agonist psychedelics (ketamine, phencyclidine, dextromethorphan, 3,4-methylenedioxymethamphetamine, scopolamine, or ibogaine)^33^ more recently than an SP. All users were excluded if they reported recent heavy cannabis use. Inclusion of cannabis users required abstinence for 4 weeks if recently using more than every other day, 2 weeks if recently using monthly to weekly, 1 week if using monthly, and 3 days otherwise—roughly corresponding to time required for cannabis-associated cognitive impairments to improve.^35^

Of the 1249 participants who completed screening, 448 were found eligible (see **Supplementary Table S1**), and 347 completed all study measures. An additional 27 participants completed at least the full first questionnaire and passed all QC checks and so were retained for final analysis pertaining to the completed data. Final analysis included 228 participants (186 SP users; **Table 1**). Eleven participants’ VCH data were excluded from analysis for failure of QC measures described above.

### Measures

#### Demographics

Participants self-reported race, age, sex assigned at birth, and psychiatric diagnoses.

#### Clinical Assessments

##### Lifetime SP-associated PPA Screener.^6^

History of SP-associated PPAs were assessed using a commonly employed^3,6,9,13,36^ 9-item questionnaire designed to evaluate unusual visual experiences beginning the day after SP use, excluding experiences better accounted for by other altered states of consciousness or pre-existing conditions. Three supplementary items were also added to capture common perceptual alterations not fully represented in the original scale: (a) seeing patterns or textures with open or closed eyes that were not actually present, (b) noticing more details in the surrounding environment, and (c) experiencing things as looking or seeming different (see **Supplementary Materials**).^37^ Participants rated both duration of their most persistent PPA and the typical intensity of PPAs when present. Duration was rated on an ordinal scale from 1 (less than 1 day) to 7 (greater than 1 year), whereas intensity was rated from 1 (seconds to minutes) to 3 (constant).

##### Cardiff Anomalous Perceptions Scale (CAPS)^38^

The CAPS queries current (past month) abnormal perceptual experiences across multiple perceptual modalities, from mild distortions to complex hallucinations. Endorsed items are followed up with Likert-scale ratings of frequency, distress, and distraction (see **Supplementary Materials**).

### Serotonergic psychedelic and other drug use

An ad hoc survey was developed to assess SP and other drug use. Participants reported days since last use, age at first use, and uses within the past six months. For participants reporting use of SPs or atypical psychedelics, the survey obtained additional information on lifetime number of uses. For SPs only, participants reported reasons for use, perceived benefit, the proportion of uses across subjective dosage categories, and participants’ preferred and most recently used SP. See **Supplementary Materials** for specific questions and dose determination guides.

#### Visual Conditioned Hallucination (VCH) Task

To assess perceptual processing, participants completed the online VCH task implemented in JavaScript via React v16.8.6 (https://react.dev/). The structure of this task is identical to other Conditioned Hallucinations tasks described previously.^26,28,29^ In the VCH task, participants performed a forced-choice visual signal detection task in which they pressed a key to report detection of a gabor patch embedded within dynamic visual white noise around a fixation cross (**Fig. 1a**). Participants rated confidence in their choice on a 1-5 scale after every trial by holding down the key signaling detection or non-detection. Participants completed two practice blocks of 10 trials each in which the gabor patch was presented only with full and absent contrast, and participants repeated these until they demonstrated at least 80% accuracy. Trials were repeated until a response was received. A pure, 500-Hz auditory tone was co-presented on every trial. Participants were instructed to foveate on the fixation cross. Gabor patch contrasts were varied via Bayesian staircasing procedure according to the Quaternion Estimator (QUEST) algorithm to determine 75% detection threshold.^39^ Contrasts corresponding to estimated 25% and 50% detection rates were then estimated from the 75% threshold and a standard psychometric curve. Gabor patch contrasts subsequently varied between 0%, 25%, 50%, and 75% estimated detection contrasts, separated into 12 blocks of 30 trials (**Fig. 1b**). Trial contrast proportions were fixed while trial order was randomized within each block. By virtue of the training blocks and a trial scheme that front-loads high-contrast trials (>85% of trials in the first block to <10% in the final six; **Fig. 1b**), the auditory tone becomes a Pavlovian conditioned stimulus wherein Gabor patch perception is the conditioned response.^28^ This robustly produces instances of target perception when no target is present, referred to here as visual conditioned hallucinations (VCHs). Previous work has demonstrated sensitivity of individuals with clinical hallucinations to conditioned hallucinations, which were accompanied by activation of sensory-specific cortical regions.^28^ From a predictive processing perspective, Pavlovian conditioning constitutes an experimentally-induced perceptual prior and peri-threshold contrast presentation provides imprecise sensory evidence, enabling behavior to reflect any bias towards prior precision in the form of VCHs.

**Figure 1.**
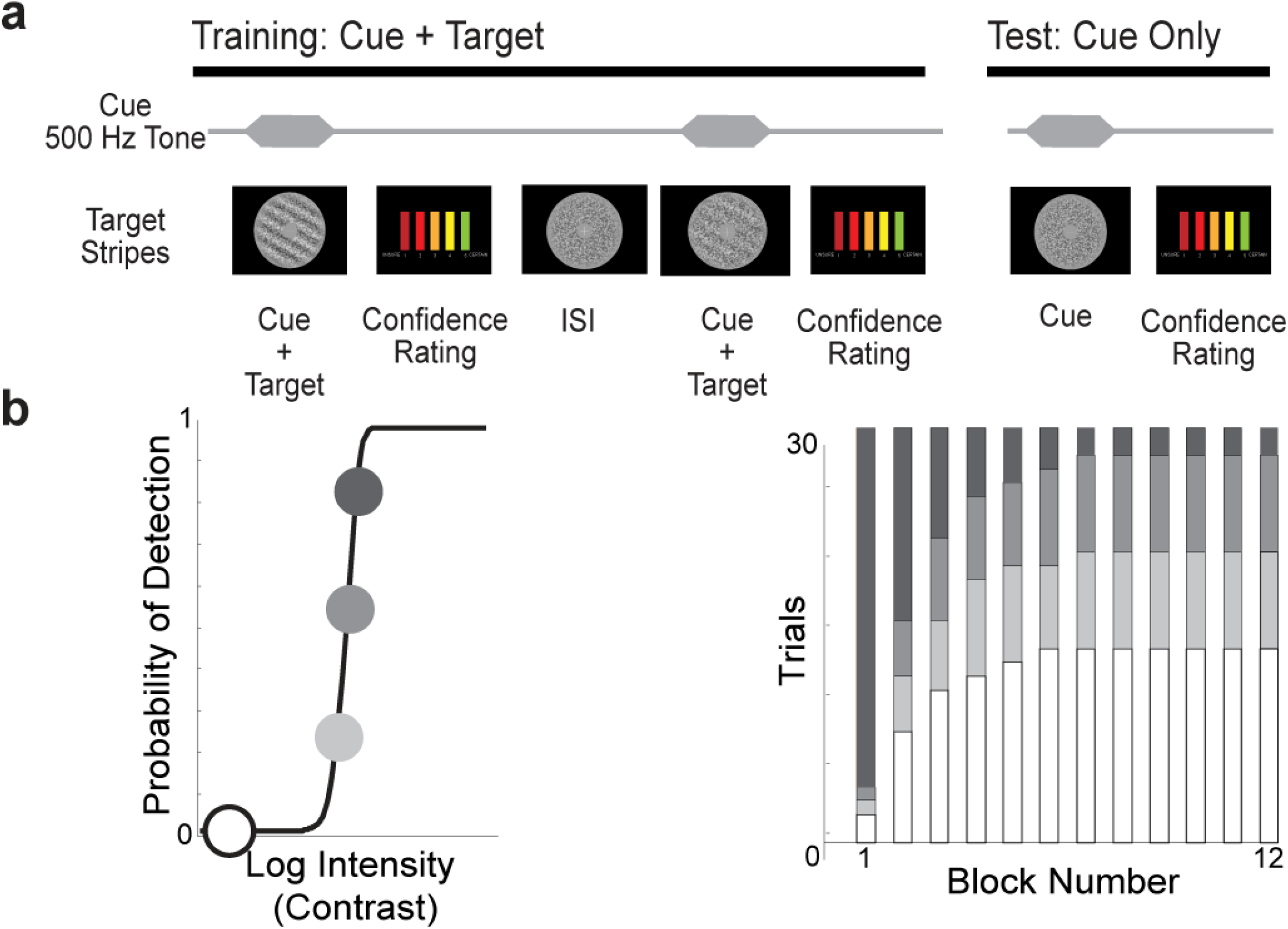
Visual Conditioned Hallucinations (VCH) task schematic. (**a**) The trial-by-trial structure in training (**left** — when some of the conditioning occurs) and the primary task when contrast intensity varies as a function of threshold (**right**). (**b**; **left**) Psychometric curve depicting (from dark to light circles) the 75%, 50%, 25%, and 0% contrast intensities determined by Quaternion Estimator (QUEST) algorithm that are assigned in each of twelve, 30-trial-blocks (**right**). Height of bars depicts the number of trials within that block of that contrast intensity.

Participants who exhibited signs of non-engagement or failed thresholding were excluded from final VCH analysis (see **Supplementary Materials**). We further considered six additional metrics of task engagement for the sake of assessing whether decreased decision precision (β) estimates were attributable to task non-engagement.

#### Hierarchical Gaussian Filter (HGF) Modeling

##### Model Description

To estimate the precision-weighting of participants’ priors in the VCH task, we employed two-level versions of the Hierarchical Gaussian Filter (HGF)^40^ as implemented in Julia (v.1.10.8)^41^ using the *HierarchicalGaussianFiltering* (v.0.6.2), *ActionModels* (v.0.6.6), and *Turing* (v.0.34.1) libraries.^42^ The HGF is a biologically plausible^43^ model of perception and learning as hierarchical Bayesian belief updating in uncertain environments, used in past work to uncover latent states driving behavior in conditioned hallucinations tasks and relating to hallucinations.^26–29^ Parameter estimates are obtained by inverting the HGF model and fitting to participant data using Bayesian inference approximated by Markov Chain Monte Carlo (MCMC) sampling, the default for the libraries listed above.

**Figure 5a** depicts the generative HGF model’s key state and trait parameters. The top tier (*X_2_*) of the HGF represents individuals’ belief in the key stimulus-outcome contingency (i.e., that the tone predicts the presence of the Gabor patch target), and the bottom tier (*X_1_*) represents this contingency in probability space (i.e. the probability of the target being present given the cue on any given trial). Beliefs at each tier are updated hierarchically in the form of precision-weighted prediction errors as inputs are observed on a trial-wise basis.

Participant choices depend on posterior belief state, μ, calculated via linear combination of the stimulus intensity *s* (based on detection thresholds above; i.e. the *likelihood*) and prediction *X_1_* (i.e. the *prior*), weighted by the parameter ν—a prior:likelihood weighting ratio. This posterior belief (μ) is converted into an action probability using a unit-square sigmoid function for which a decision precision (β) session parameter governs the steepness (**Supplementary Equation 1**), effectively capturing how deterministic participants are in their response, given their posterior percept (**Supplementary Fig. S1**). Participant choices are then sampled from this final action probability distribution. One final session parameter—the evolution rate or volatility (ω) of *X_2_*— is also estimated.

We considered two additional variants of this HGF: 1) adding a third level (*X_3_*) that encodes beliefs about the phasic volatility of *X_2_* as well as a meta-volatility ω*_2_* evolution rate parameter, 2) using empirically-derived detection probabilities (i.e. 25%, 50%, 75% → 41.80%, 71.15%, 89.94%) for *s* from a separate sample^26^ of participants endorsing no history of visual or auditory hallucinations (n = 29); this latter model was tested because detection probabilities were higher than those estimated by QUEST, suggesting that sensory evidence may have been more precise than expected (see **Supplementary Materials** and **Supplementary Fig. S2**). This resulted in a total of four models examined in model comparison. Please see **Supplementary Fig. S3** for a full description of the Bayesian workflow for computational psychiatry including model comparison.^44^

### Parameter Estimation, Model Comparison, and Model Validation

We followed the recently described Bayesian workflow for generative computational modeling in psychiatry.^44^ The 2-level model with empirically-derived detection probabilities used for presented analyses was selected by Random-effects Bayesian model selection (RFX-BMS) with a protected exceedance probability (PXP) of 1.00 and a posterior frequency (Ef) of 0.803. This model demonstrated excellent parameter recovery for ν, β, and ω (0.940, 0.837, and 0.784). Further details are described in the **Supplementary Materials**.

### Signal Detection Analysis

To supplement the above measures and other behavioral variables from the VCH task, we calculated two key Signal Detection Theory parameters: d’ (d-prime) and Criterion (c). D’ represents a participant’s ability to discriminate signal from noise and is calculated as

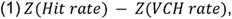

Where Z() represents the inverse cumulative distribution of the normal function, hit rate is the number of hits divided by the number of target-present trials and VCH rate is the number of VCHs out of total stimulus absent trials. A log-linear correction was applied to account for participants without any VCH trials.^45^ Criterion represents participants’ response bias or threshold—the stimulus strength after which a participant reports detection (either as a “hit” or VCH). It is calculated as:

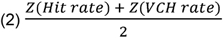

Hence, a Criterion value below zero indicates a more liberal response style that requires less intense stimuli to report detection, while a value above zero represents the opposite.

### Statistical Analyses

#### General Analytic Approach

As a first-pass assessment of variables associated with current PPA symptoms and PPA history, we first used non-parametric Spearman’s ρ (for the continuous CAPS visual items score) or Mann-Whitney-U tests (for binary PPA history). We estimated 94% confidence intervals for mean differences in HGF belief trajectories using bootstrap sampling with replacement for 10,000 iterations. We more rigorously investigated variables showing signs of an association, accounting for variable magnitude and potential covariates, using linear, Bayesian regression models (BRMs) for their flexibility in accommodating unusually distributed response variables and easy translation into interpretable mediation hypothesis testing.

All values following a “±” symbol represent SD unless otherwise noted. All frequentist tests were two-tailed.

In agreement with the American Statistical Association’s statement on p-values and significance testing,^46^ we do not use any arbitrary threshold value for posterior probability or p-values, but rather consider them continuously alongside effect sizes. Due to the exploratory nature of all analyses, we also do not correct for multiple tests.

All visualizations and figures were created using matplotlib version 3.10.7 (https://matplotlib.org/stable/install/index.html) and seaborn version 0.13.2 (https://seaborn.pydata.org/) in Python v.3.12.4 (https://www.python.org/downloads/).

#### Bayesian regression model diagnostics

BRMs were implemented using the *BRMS* package (v. 2.23.0)^47^ in R version 4.4.1^48^ and RStudio version 2024.12.1.563^49^ with default priors (including flat priors for slope), 4,000 iterations (+ 6000 warmup), and four chains. BRM likelihood functions were selected based on the empiric distribution of each dependent variable and verified based on visual inspection of posterior predictive checks and residual patterns (see below). All presented BRMs had R (the Gelman-Rubin statistic for confirming chain convergence) values within 1 ± 0.01, effective sample sizes over 1000, and no divergent transitions after startup.

We confirmed basic model fit for all BRMs via visual inspection of posterior predictive plots and simulation-based residual diagnostics implemented within the *DHARMa* package^50^ (see **Supplementary Materials & Fig. S4**).

#### Mediation Analysis

To test whether identified PPA correlates were specific to *SP-related* PPAs as opposed to correlates of PPAs in general, we tested whether any of VCH/HGF-derived variable mediated observed associations between SP use patterns and PPAs. We examined only those SP use patterns associated specifically with SP-related PPA history and current CAPS visual item endorsements—age of first SP use and average SP dose, respectively. To infer mediation, we leveraged a similar framework in multivariate models in which we simultaneously fit two models:

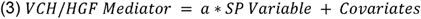

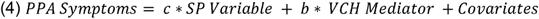

Here, the indirect or mediated effect is calculated by sampling 1000 predicted mediator values from the model described in **equation (3)** for mean and +1 SD predictor values for all 16,000 joint parameter draws, and then sampling predicted response variable values from the model described by **equation (4)** for each.^51,52^ The proportion of simulations suggesting a change in the PPA symptoms is the probability of an effect; see **Supplementary Materials** for more information on specification and inference method from our Bayesian regression models.

For **equation 3** above, where we model VCH and HGF variables, we specify “Student *t*” likelihoods as outlier-robust alternatives to roughly-gaussian distributed variables^53^ for all but two variables: 1) VCH rate, for which we specify a zero-inflated beta distribution,^54^ and 2) the ν (prior-weighting) parameter, for which we specify a gamma distribution.

### Covariates and sensitivity analyses

For primary analyses presented here, we covaried for age, sex assigned at birth, IQ approximated using a 9-item version of Raven’s progressive matrices,^55^ and mental illness history. We also conducted sensitivity analyses (summarized in **Supplementary Fig. S4** and **S5**) with varying covariates and data subsets including: 1) adding 42 SP-naïve participants, 2) removing outlier observations more than 150% the IQR of the predictor variable, 3) controlling for clinical/demographic variables with some evidence of association with PPA history or CAPS vision items, 4) only controlling for age, and 5) no covariates. For models predicting CAPS visual endorsements only, we also conducted sensitivity analyses that added 42 SP-naïve individuals. For models predicting SP-related PPA history only, we conducted sensitivity analysis excluding those with missing CAPS data to ensure that results did not vary by sample. Notably, 56 (40 SP users) and 16 (9 SP users) participants failed to complete the CAPS and VCH task, respectively (see **Supplementary Materials**) and were excluded listwise from models using these variables.

Additional covariates used in sensitivity analyses were selected if they appeared to systematically vary between participants with and without SP-associated PPA history and CAPS vision endorsement. This was determined via chi-square tests of independence; we selected variables with p values close to 0.1 and sufficient observations at different variable levels to be used in regression analysis (see **Table 1**). Because CAPS visual items are semicontinuous, we also selected variables that exhibited signs of association as assessed by spearman correlations (**Supplementary Table S2**).

For all sensitivity analyses, we note significant (>15%) changes in inferred effect probability in the text. Otherwise results were robust to sensitivity analyses.

### AI Use Disclosure

Claude Sonnet and Opus 4.6 as implemented in Claude Code were used for programming including data analysis, figure creation/assembly, the Bayesian workflow for generative computational modeling in psychiatry, as well as maintaining documentation in our available code. The initial code-base for statistical tests were hand-coded, but all tests were touched by an AI agent and not all lines of code have been read by a human. All outputs were, however, subject to rigorous verification for validity—varying by application, but including verifying formulas, variable normalization, visualizations, and other expected statistical behavior. All **Results** figures and statistics as well as plots and statistics in the **Supplementary Materials** were created using AI as a tool.

## Results

### Demographics, clinical, and SP use history

The sample skewed young, male, white, United-States (US)-residing, and well-educated, with more than 59% completing a four-year degree or greater (**Table 1**). There were virtually no differences in demographic profile between participants endorsing a history of SP-associated PPAs or current, CAPS-assessed PPAs. Those with a SP-associated PPA history were less likely to be multiracial, and those with current PPAs were younger (consistent with other reports),^3^ more likely to be European, and less likely to be US-residing. Psychiatric illness rates were similar to the general population.^56^ As expected, polydrug use was considerable; all participants had used another psychoactive drug and the majority had used one in the past month. Most participants reported prior non-serotonergic psychedelic use, though fewer than 10% had used in the past month.

SP use patterns are summarized in **Supplementary Figure S6**. Total lifetime uses varied significantly (44.7 ± 94.3) and were positively skewed—from 1 to 700. The weighted average subjective SP dose used was more consistent (252.2 μg ± 175.1 μg), with the vast majority of uses concentrated at the highest end of doses administered in contemporary in-laboratory research (the equivalent of ∼200μg of LSD).^57^ More than a quarter of participants reported doses substantially higher than this cutoff. First SP use generally occurred in young adulthood (22.9 ± 8.9 years), with 73% of participants reporting initiation prior to age 25 and only 9% at age 40 or older. The plurality of participants reported using SPs for both therapeutic and recreational reasons, whereas the remainder were split relatively evenly between primarily recreational and primarily therapeutic use (see **Supplementary Fig. S6**).

### PPA History & Current PPAs

Consistent with prior findings,^6,7,10^ SP-associated PPAs were common, endorsed by 131 (57.5%) of the full sample (**Fig. 2**). Also consistent,^3,8,9^ PPAs typically occurred after few SP uses (median 3; IQR [1,5]). Thirty-three participants (25.6%) reported effects after just one dose. Participants generally endorsed multiple PPA phenomena; the modal symptom count was 3 (17.6%), and 78 (59.5%) endorsed 4 or more symptoms. The most endorsed PPA was “greater color intensity”, reported by 28 (21.4%). Also consistent with other studies, most (81; 61.8%) reported that symptoms did not persist beyond 1 week, whereas 11 (8.4%) reported durations longer than 1 year. Finally, as reported before,^6–9^ the vast majority of participants reported not experiencing distress (121; 92.4%) nor seeking treatment (129; 98.5%) and only 3 (1.3%) reported a formal HPPD diagnosis.

**Figure 2.**
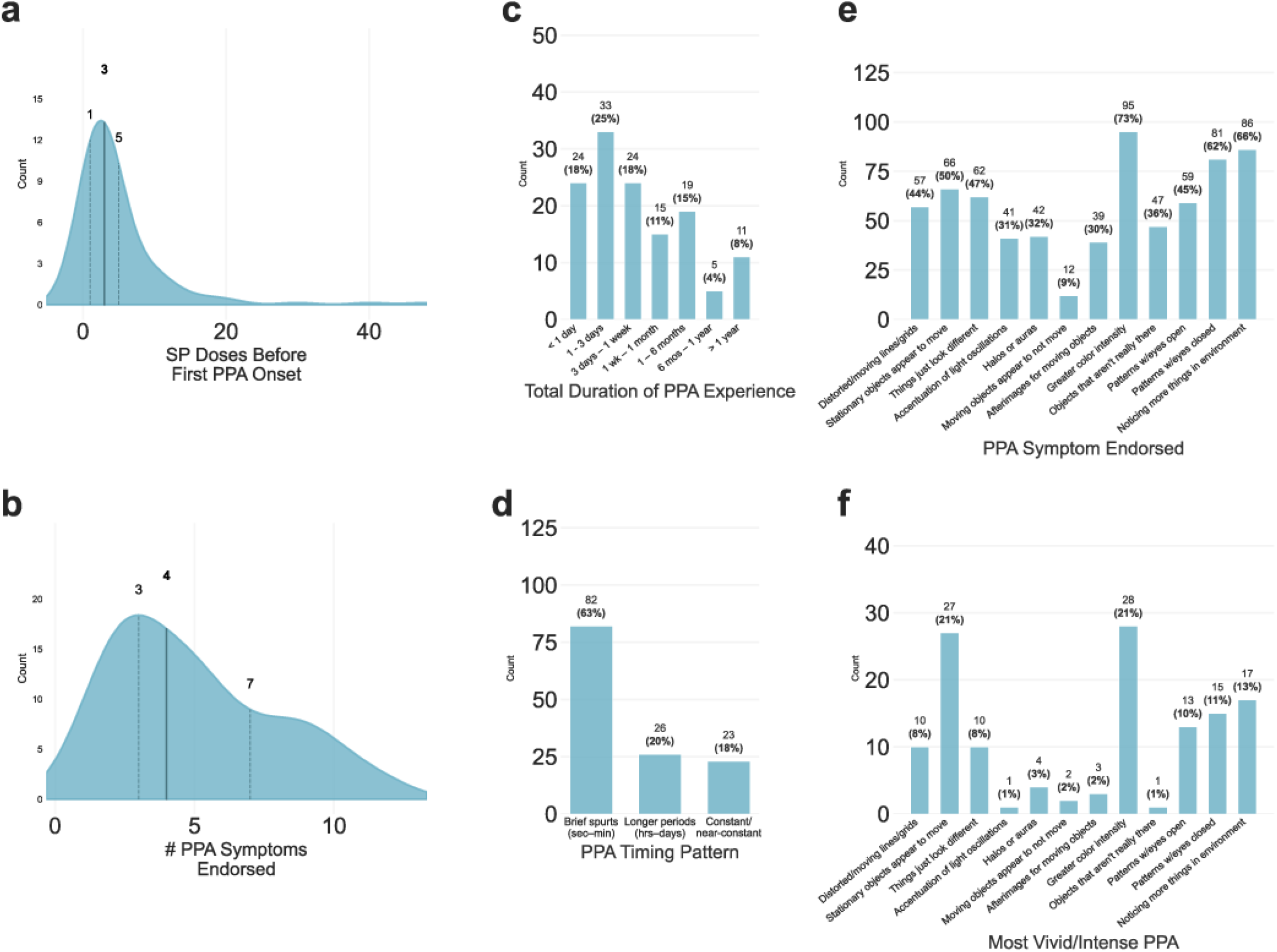
Characteristics of SP-associated PPAs. Density plots display the distributions of (**a**) how many times participants had taken SP macrodoses before first noticing a PPA and (**b**), number of different PPAs participants had experienced after SP use. Dark lines and bolded values indicate medians while dotted lines and non-bolded labels indicate 25th and 75th percentile values. Bar plots qualify SP-associated PPAs by (**c**) the longest any PPA ever lasted (from less than 24 hours to greater than 1 year), (**d**) the most typical chronologic character of PPAs when active (from “flashback”-style effects lasting less than a few minutes to a constant shift in perception as in certain HPPD subtypes) across all PPAs, (**e**) which PPA phenomena were ever experienced after SP use, as well as (**f**) which phenomenon was the most “vivid/intense.” See **Supplementary Materials** for SP-associated PPA screening questionnaire and follow-ups. (n = 131)

The majority (52.3%) of SP users with CAPS data endorsed one or more current PPAs—primarily those endorsing prior SP-associated PPAs (86.8%). Consistent with SP-associated PPAs, the most endorsed CAPS item was “lights seeming brighter or colors seeming more intense”, and this effect was typically rated as non-distressing and non-distracting, but also one of the most frequent symptoms (**Fig. 3**). Frequency was episodic, with only 13.2% endorsing any symptoms “all the time”, and most symptoms experienced “not often” to “sometimes”. Distress was rarer, with 47.1% of participants reporting one or more PPAs endorsing all of their experiences as “not at all” distressing, while only 11.8% endorsed an experience as “firmly” or “very” distressing.

**Figure 3.**
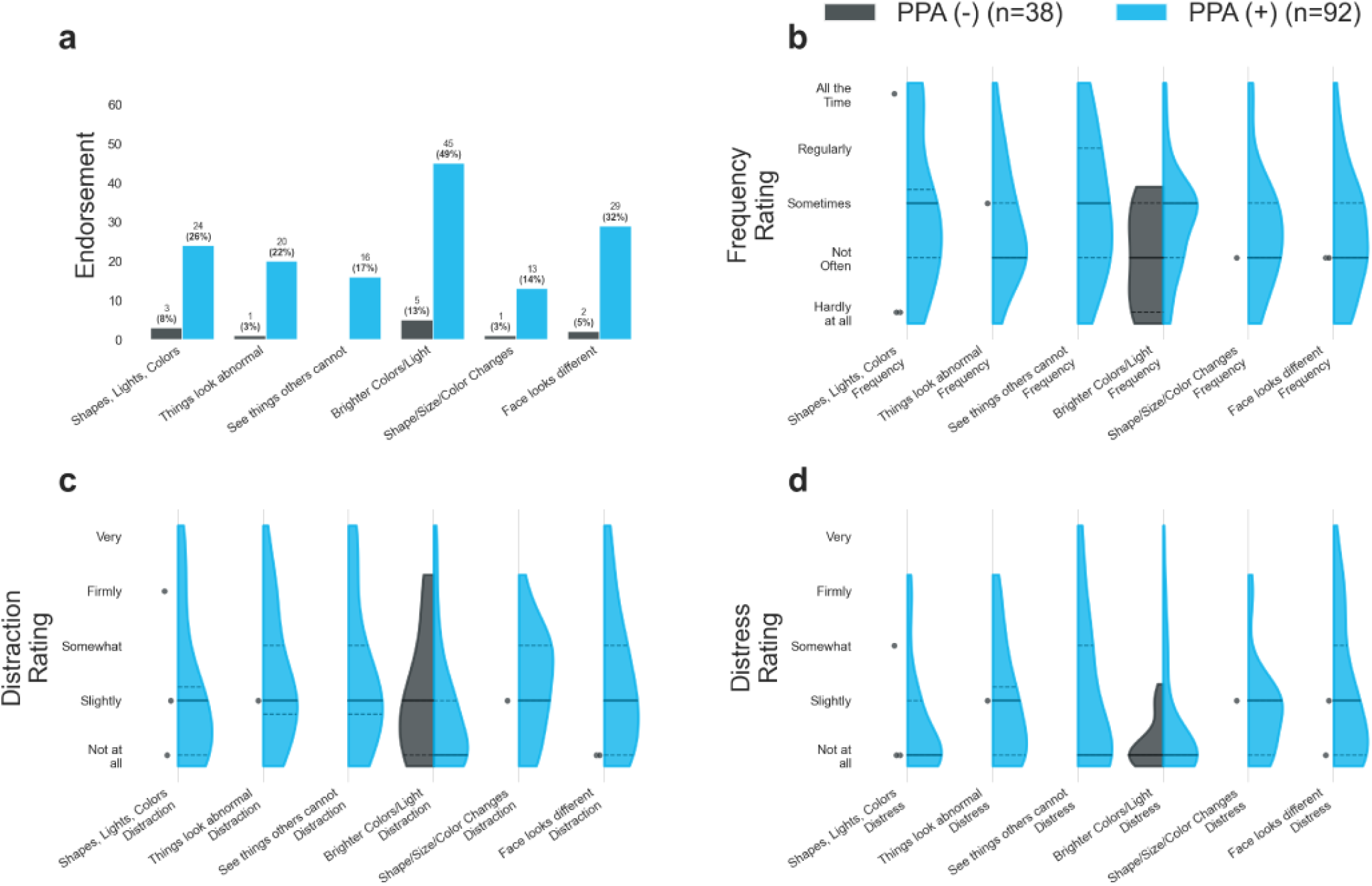
Phenomenology of CAPS-Assessed visual PPAs in the past month and their severity. (**a**) Endorsement rates for each visual symptom on the CAPS split by whether participants had endorsed SP-associated PPAs on a 12-item screener—see **Supplementary Materials** for CAPS vision items and 12-item screener. For those endorsing each CAPS item in (**a**) violin plots display participants’ subjective rating of that item in terms of its (**b**) frequency—from “Hardly at all” to “All of the time”, (**c**) level of distraction—from “Not at all” to “Very” and (**d**) distress caused—from “Not at all” to “Very”. Solid lines indicate median values, and dotted lines indicate IQR values in violin plots. Electric blue denotes SP-associated PPA endorsement and dark gray denotes denial. (n = 130; 92 endorsing SP-associated PPAs)

### Earlier age at first SP use is associated with a greater lifetime risk of SP-associated PPAs

We first examined which patterns of SP use were associated with PPA history among SP users. Compared with those who denied a history of PPAs after SP use, participants who endorsed PPAs (PPA+) had a younger age at first SP use (**Fig. 4a**; U = 4624.5, p = 0.002; see **Supplementary Table S3** for additional statistics from all binary Mann-Whitney U tests). By contrast, there was less certain evidence that PPA+ individuals have had greater lifetime SP uses (U = 3040, p = 0.093) and no credible sign of greater average doses used (U = 3238, p = 0.277). We further interrogated this association with a Bayesian regression incorporating relevant covariates (age, sex, IQ, and mental illness history) and magnitude of both variables; we found that younger (-8.9 years) age of first SP use was associated with a 12.0% higher estimated probability of lifetime PPAs (Δ = - 12.0%, P(Δ<0) = 99.9%, 94% HDI [-20.3%, -4.2%]). Greater lifetime SP uses (94.3 uses) were associated with greater PPA probability (Δ = 9.5%, P(Δ>0) = 98.7%, 94% HDI [1.3%, 17.8%]) but this effect diminished considerably when excluding outliers (Δ = 13.4%, P(Δ>0) = 83.3%, 94% HDI [-12.6%, 23.8%]; 10.8% of participants dropped; **Supplementary Fig. S4**). Higher average SP dose used (+175.1 μg) was not estimated to credibly influence PPA probability (Δ = 1.0%, P(Δ>0) = 60.9%, 94% HDI [-5.5%, 7.7%]); though this changed after outlier exclusion (Δ = 11.4%, P(Δ>0) = 99.5%, 94% HDI [3.9%, 18.4%]; 5.9% of participants dropped). In summary, only younger age of first SP use was robustly associated with small but meaningful increases in probability of prior SP-associated PPAs, and so subsequent mediation analyses examining past PPA risk focus on this predictor.

**Figure 4.**
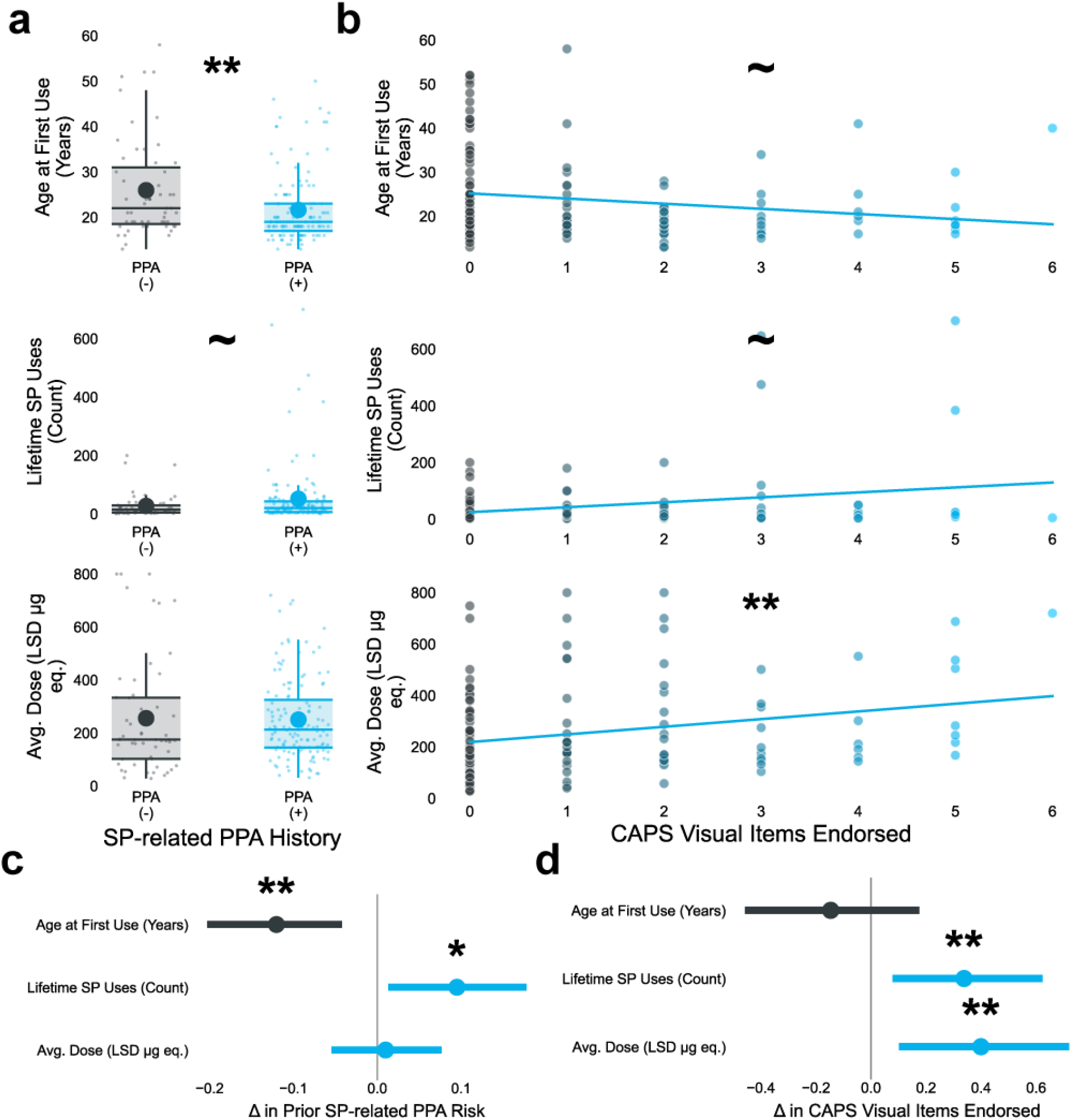
SP-use patterns associated with prior and current PPAs. Differences in (**top**) age of first SP use, (**middle**) total lifetime SP uses, and (**bottom**) average SP dose used by (**a**) endorsement of any prior SP-associated PPA assessed (n=186) and (**b**) number of past-month visual CAPS items endorsed (n=130). Boxplots’ horizontal lines indicate (from bottom-to-top) 25%, 50%, and 75% percentiles; vertical lines indicate 150% of the interquartile range; dark colored dots indicate means; and asterices indicate p-values from Mann-Whitney U-tests. Scatterplots’ lines represent a linear regression line, and asterices indicate p-values from spearman correlations. Dark gray to electric blue indicates more PPA burden. Forest plots depict 94% Highest Density Intervals (HDIs) from covariate-adjusted Bayesian regression models (BRMs) expected change in (**c**) probability of prior SP-associated PPAs and (**d**) number of past-month visual CAPS items endorsed from a 1 SD increase in SP use variables. Dark gray indicates that changes are expected to cause a reduction in PPA risk or CAPS symptoms while electric blue indicates an increase. (∼p <0.1, *p □ < □ 0.05, **p □ < □ 0.01, ***p □ < □ 0.001).

### Higher average SP doses are associated with more current PPAs

We next asked how these SP patterns varied with current (past-month) visual PPAs (**Fig. 4b**). We controlled for age because age correlated with all SP use variables and strongly anticorrelated with CAPS endorsements (**Table 1**). Average SP dose (ρ = 0.249, p = 0.004) strongly correlated with greater CAPS vision items endorsed, whereas age of first SP use (ρ = -0.156, p = 0.078) and lifetime SP uses (ρ = 0.148, p = 0.094) had 50% weaker correlations. Regression models confirmed that a higher average dose was associated with a greater estimated number of CAPS vision items endorsed (Δ = 0.399, P(Δ>0) = 99.5%, 94% HDI [0.101, 0.720]). Age of first SP use showed no association with CAPS vision (Δ = -0.146, P(Δ<0) = 81.1%, 94% HDI [-0.458, 0.176]). Lifetime SP uses were estimated to increase CAPS vision items (Δ = 0.338, P(Δ>0) = 99.8%, 94% HDI [0.078, 0.623]), but this effect was entirely outlier-driven (Δ = 0.066, P(Δ>0) = 53.7%, 94% HDI [- 0.753, 3.482]). Subsequent mediation analyses of current PPA symptoms focus on higher average SP doses used, as this was the clearest predictor of current PPA symptoms.

### Lower visual detection thresholds and higher VCH Rates are associated with both PPA history and current CAPS visual symptoms

We next investigated potential behavioral correlates of SP-associated PPA history and current PPAs. Those with a history of SP-associated PPAs had lower detection thresholds (i.e., reported detection at lower visual contrasts; U = 3206, p = 0.028; **Fig. 5a**), associated with a 10.3% increase in PPA risk (P(Δ<0) = 99.5%, 94% HDI [-18.6%, -2.3%]; **Fig. 5c**). PPA (+) participants also had greater VCH rates (U = 1948, p = 0.011), associated with an estimated 8.3% increase in PPA risk (P(Δ>0) = 98.6%, 94% HDI [1.4%, 15.0%]). Veridical detection in the easiest-to-detect 75% condition (hit-rate) was not related to PPA history (U = 2596.5, p = 0.925; Δ = -1.6%, P(Δ<0) = 67.2%, 94% HDI [-8.7%, 5.1%]).

**Figure 5.**
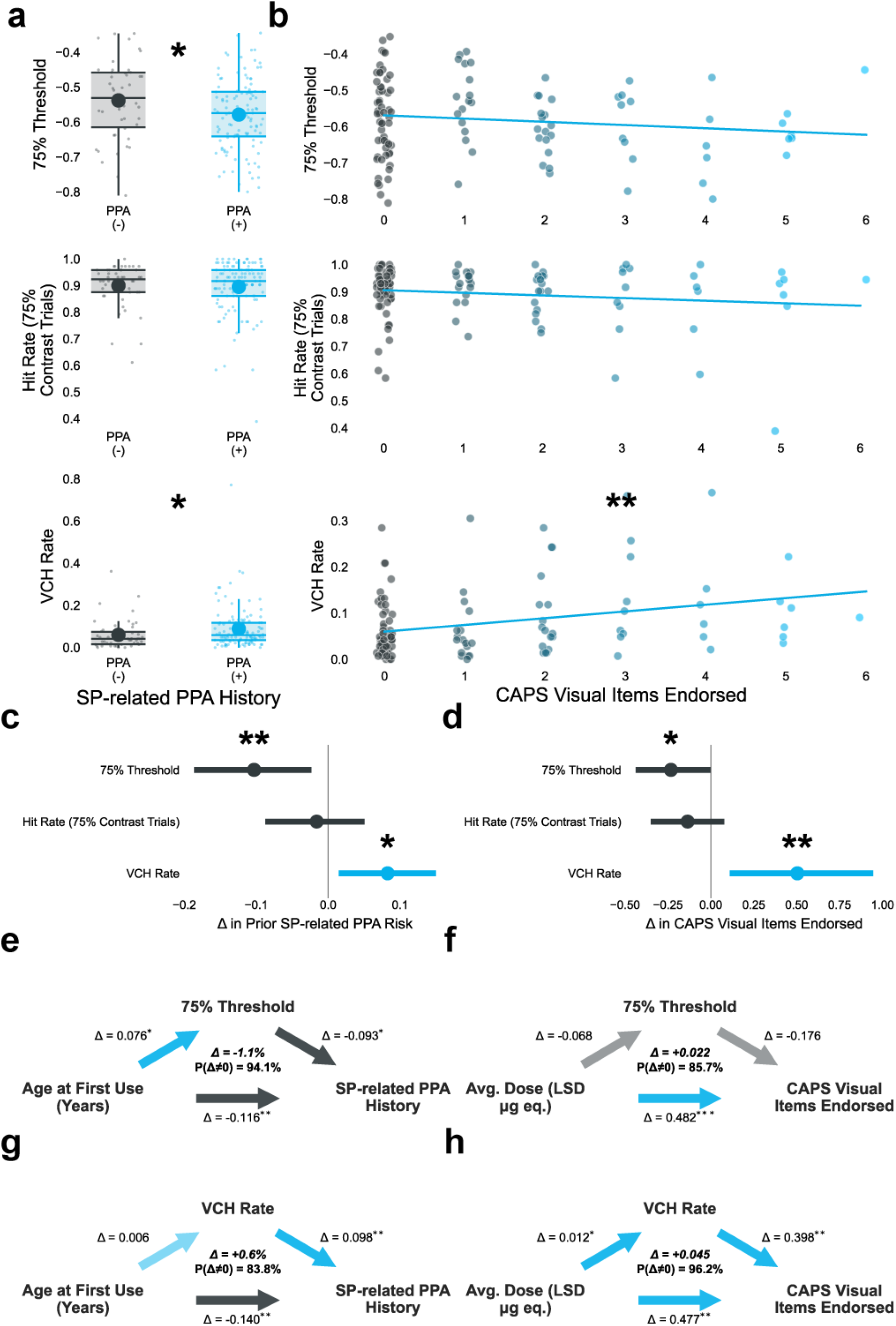
VCH task behavioral measures associated with prior and current PPAs. Differences in (**top**) QUEST-estimated 75% detection threshold for gabor patch contrast intensity, (**middle**) correct detection

CAPS items showed a weaker correlation with threshold (ρ = -0.103, p = 0.279) that strengthened after controlling for age (ρ__partial_ = -0.153, p = 0.108; **Fig. 5b**). Lower (-0.105) thresholds were associated with a 0.233 increase in CAPS vision endorsements (P(Δ<0) = 96.2%, 94% HDI [-0.439, -0.002]; **Fig. 5d**). Higher VCH rates (ρ = 0.304, p = 0.001) correlated with current CAPS visual items endorsed and was associated with 0.504 more CAPS visual items (P(Δ>0) = 99.6%, 94% HDI [0.110, 0.946]). Hit rate was not correlated with CAPS items (ρ = -0.040, p = 0.677), though it exhibited uncertain evidence of decreasing CAPS endorsements (Δ = -0.135, P(Δ<0) = 89%, 94% HDI [-0.349, 0.079]).

We then asked if greater visual sensitivity (lower thresholds) and higher VCH rates mediated the SP use patterns previously associated with past and current PPAs. The relationship between younger age of first use and increased PPA history probability was plausibly mediated by lower visual threshold (Δ = -1.1%, P(Δ<0) = 94.1%, 94% HDI [-3.3%, 0.3%]; **Fig. 5e**), but not VCH rate (Δ = 0.6%, P(Δ>0) = 83.8%, 94% HDI [-0.7%, 2.1%]; **Fig. 5g**). By contrast, the relationship between SP dose and current PPAs was not confidently mediated by threshold (Δ = 0.022, P(Δ>0) = 85.7%, 94% HDI [-0.021, 0.106]; **Fig. 5f**), but showed modest evidence of mediation by VCH rate (Δ = 0.045, P(Δ>0) = 96.2%, 94% HDI [-0.012, 0.139]; **Fig. 5h**).

### Decreased decision precision is predictive of both prior SP-associated PPA risk and current PPAs, and increased prior weighting may be predictive of current PPAs

Finally, to determine which latent states drove these behavioral patterns, we fit HGF models to VCH behavioral data. We then examined whether HGF-derived parameter estimates varied with both PPA measures.

Those with a history of SP-associated PPAs exhibited no clear difference in estimated prior weighting, ν (U = 2607, p = 0.956), and ν did not increase PPA history probability (Δ = 1.6%, P(Δ>0) = 68.7%, 94% HDI [-4.7%, 7.7%]; **Fig. 6h**). By contrast, PPA (+) participants did have more stochasticity in responding, reflected in lower β values (U = 3226, p = 0.023; **Fig. 6j**). Lower β was associated with a 7.7% increased probability of prior SP-associated PPAs (P(Δ<0) = 98.1%, 94% HDI [-15.6%, -0.9%]; **Fig. 6l**). There were no associations between PPA risk and ω (U = 2793, p = 0.520; Δ = -0.9%, P(Δ<0) = 60%, 94% HDI [-8.1%, 5.4%]; **Fig. 6b**) nor belief trajectories (Δ = 0.094, 94% CI [-0.092, 0.304]; **Fig. 6d** & **f**).

In the case of current PPAs, β again negatively correlated with endorsements (ρ = -0.225, p = 0.017; **Fig. 6k**), and was associated with a -0.292 reduction in current PPAs (P(Δ<0) = 99.2%, 94% HDI [-0.499, -0.086]; **Fig. 6m**). We also saw a very weak association with prior weighting, ν (ρ = 0.139, p = 0.143). Prior weighting was estimated to increase visual PPAs by 0.259 with modest confidence (P(Δ>0) = 96.4%, 94% HDI [-0.030, 0.588]; **Fig. 6i**). This effect significantly diminished with outlier exclusion (Δ = 0.128, P(Δ>0) = 74.9%, 94% HDI [-0.233, 0.600]; 2.7% of participants dropped) and remained uncertain even after controlling for β (Δ = 0.192, P(Δ>0) = 83.4%, 94% HDI [-0.217, 0.724]; **Supplementary Fig. S4**). Current PPAs were also not associated with ω (ρ = -0.068, p = 0.476; Δ = -0.012; P(Δ<0) = 53.1%, 94% HDI [-0.297, 0.294]; **Fig. 6c** & **m**) nor belief trajectories (Δ = 0.168, 94% CI [-0.028, 0.365]; **Fig. 6e** & **g**).

β did not mediate the increased PPA risk from younger age of first SP use (Δ = 0.2%, P(Δ>0) = 62.9%, 94% HDI [-1.2%, 1.8%]; **Fig. 6n**), apparently because age of first use did not influence β (Δ = -0.016, P(Δ<0) = 63.9%, 94% HDI [-0.097, 0.073]). However, the relationship between average SP dose and increased CAPS visual item endorsements was plausibly mediated by decreased β (Δ = 0.058, P(Δ>0) = 95.7%, 94% HDI [-0.015, 0.166]; **Fig. 6o**). Unsurprisingly, ν did not mediate age of first use effects on PPA history (Δ = 0.1%, P(Δ>0) = 64.3%, 94% HDI [-0.8%, 1.3%]) nor average dose effects on current PPAs (Δ = 0.018, P(Δ>0) = 81.1%, 94% HDI [-0.038, 0.138]; **Supplementary Table S5**).

### Decreased decision precision, PPA history, and current PPAs are related to diminished criterion, signal-versus-noise discriminability, and more confident VCHs

From the results above, low decision precision (β) appeared to be the strongest explanatory parameter. However, its interpretation is ambiguous: low decision precision could reflect poor task engagement, impaired sensory processing, and/or altered metacognitive processing. To help determine which of these interpretations is best supported by the data, we conducted a series of exploratory analyses.

We first considered whether lower β scores indicated noisy responses due to poor task engagement. We examined correlations between β and six separate measures of task engagement, including repeated responses, response randomness, and response time-based measures(see **Methods** and **Supplementary Fig. S7**) and found no evidence that poor task engagement related to low decision precision.

We next examined participants’ overall task performance, splitting participants by the median values of the two PPA-linked HGF parameters, β and ν (**Fig. 7a**). While both were associated with higher VCH rates, lower hit rates in the target-present portion of the task drove lower β estimates, and those with the greatest VCH rates also had lower hit rates, resulting in flatter psychometric curves, classically corresponding to low sensitivity^58^. Consistent with this, we found direct correlations (**Fig. 7b**) between β and both d (ρ = 0.519, p < 0.001) and decision criterion (ρ = 0.433, p < 0.001)—suggesting poor sensory fidelity alongside a low threshold for target perception. Given associations between lower threshold and SP-related PPAs above, we also considered whether diminished hit rates associated with low β were due simply to the task being systematically more challenging for those with lower threshold. After controlling for age, threshold exhibited a very uncertain correlation with hit rate (ρ = 0.124, p = 0.120) and no correlation with β (ρ = 0.004, p = 0.956). Threshold more strongly correlated with d (ρ = 0.146, p = 0.067). These results suggest that threshold alone does not explain beta’s association with poor sensory sensitivity.

**Fig. 7.**
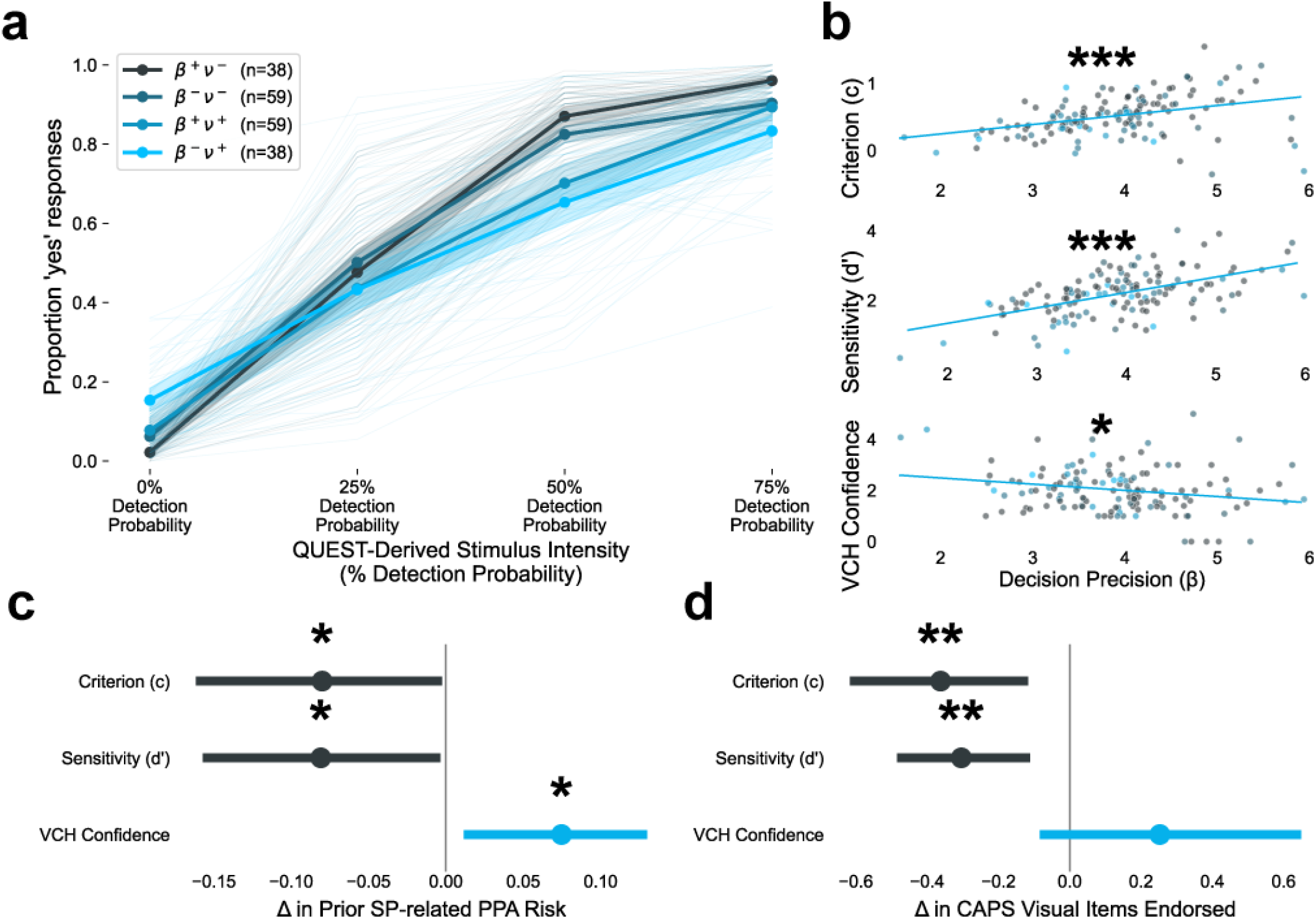
Behaviors associated with low decision precision (β) and their association with PPAs. (**a**) Psychometric curves split based on relative β and ν estimates, splitting at their median value. β*^+/-^ =* above/below median β value; ν^+/-^ = above/below median ν value. (**b**) Spearman correlations between β estimates and d, Criterion, and confidence on VCH trials. Electric blue represents participants with more CAPS visual item endorsements while dark gray represents fewer. Forest plots depicting 94% HDIs from covariate-adjusted Bayesian regression models (BRMs) expected change in (**c**) probability of prior SP-associated PPAs and (**d**) number of past-month visual PPAs on the CAPS from 1 SD increases in d, Criterion, and confidence on VCH trials. (∼p <0.1, *p□ < □ 0.05, **p □< □ 0.01, ***p □ < □0.001).

Finally, we considered whether there was evidence of *metacognitive* abnormalities by inspecting confidence ratings: because β is outside the HGF’s perceptual model, low decision precision could reflect lower confidence in the posterior derived from the perceptual model. We found the opposite: lower β correlated with greater confidence in VCHs (ρ = -0.194, p = 0.023; **Fig. 7b**).

Having found evidence that lower β is associated with poor sensory discrimination coincident with a liberal criterion for signal detection and reduced metacognitive accuracy, we asked whether these behaviors were also linked to PPAs. Greater PPA history probability (**Fig. 7c**) was associated with lower d (Δ = -8.1%, P(Δ<0) = 98.3%, 94% HDI [-15.7%, -0.3%]), lower decision criterion (Δ = -8.0%, P(Δ<0) = 97.9%, 94% HDI [-16.1%, -0.3%]), and higher VCH confidence (Δ = 7.5%, P(Δ>0) = 98.3%, 94% HDI [1.2%, 13.0%]), while greater current PPA count (**Fig. 7d**) was confidently associated with lower d (Δ = -0.304, P(Δ<0) = 99.7%, 94% HDI [-0.485, -0.111]) and decision criterion (Δ = -0.362, P(Δ<0) = 99.2%, 94% HDI [-0.616, -0.116]). It was less confidently associated with VCH confidence (Δ = 0.253, P(Δ>0) = 92.8%, 94% HDI [-0.084, 0.649]).

## Discussion

This study sought to identify clinical characteristics, visual behavior and computational estimates of perceptual inference associated with SP-related PPAs. Our study was designed to determine whether SP-related PPAs result from persisting prior *hypo-*precision as described in recent theories of SP action or prior *hyper-*precision as seen in psychosis-spectrum populations.^26–29^ Our results do not perfectly fit either prediction. SP-associated PPAs and current PPA burden were associated with greater VCH rates, which have been linked to an over-reliance on priors (**Fig. 5c** - **d**). We did not see consistent, parallel associations with prior hyper-precision (ν) (**Fig. 6h** - **i**), although this may be supported by consistent biases towards detection in PPAs, manifested as lower criterion estimates and lower yes/no thresholds. Modeling choices may also have led to an under-estimation of this parameter (**Supplementary Table S5**). Instead, we found that lower decision precision was reliably associated with both past (**Fig. 6l**) and current PPAs (**Fig. 6m**) and mediated the relationship between higher SP doses and current PPAs (**Fig. 6o**). Post-hoc analyses tied this lower decision precision to several elements of impaired sensory processing (**Fig. 7**). Together, this points to a potentially minor role for prior hyper-precision in the setting of disrupted sensory processing as being a primary driver of persisting perceptual abnormalities.

In this sample, we both replicate and identify novel clinical observations associated with SP-related PPAs. Consistent with prior reports, PPAs were common, rarely distressing, and typically lasted for brief periods, although sometimes extended for months to years (**Fig. 2**)^6,7,10^ Replicating prospective cohort studies’ findings,^3,9^ younger age appeared to confer special risk. Unlike prior research,^9^ SP dose was the most consistent predictor of current PPAs (although this discrepancy may be related to the high doses used and the manner of dose measurement in this study), and no associations with psychiatric diagnosis^9,59^ or gender^9^ were found. Because both dose^60^ and youth^61^ are predictive of more intense psychoactive effects, intensity of acute experiences may be a useful predictor of subsequent PPA risk.

The combination of poor sensory discriminability, low criterion, and high VCH rates and confidence (**Fig. 7c** - **d**) suggest that PPAs may result from a noisy visual system biased toward detection in the absence of appropriate metacognitive checks. This is most readily evinced in the disjunction between low estimated QUEST-derived thresholds and subsequent inaccurate detection performance in those with PPAs, indicating that low threshold is likely driven not by sensory sensitivity but a bias toward detection. Combined with low decision precision and equivocal elevations in relative prior precision, this nuanced picture does not conform to the typical binary framing of relative prior and sensory precision. Rather, it implies a dual role for bottom-up noise–simultaneously degrading ascending sensory information and acting as evidence for the presence of an expected signal despite its absence. Recent accounts of the instantiation of predictive processing within sensory cortical columns is consistent with this possibility: the expected reception of a specific input may act as a template for that input, and shape incoming sensory information to that expectation, enhancing congruent and suppressing incongruent information.^62–64^ In the setting of driving sensory noise, the template itself may produce the expected percept.^65–67^ Thus, expectations of a sensory event in the context of driving, noisy sensory information may result in a perceptual manifestation of that event. Within the context of our model, because both sensory evidence and priors exhibit elevated precision, the relative prior precision term would exhibit only equivocal elevations, as we observe here. Similar explanations have been invoked to explain HPPD and the highly comorbid visual snow syndrome^37,68^.

This computational framework may also help explain some phenomenological aspects of the PPAs reported here (**Fig. 2**): rather than true hallucinations, the most commonly reported PPAs here and elsewhere^3,9^ historically^37^ have been enhancements of color or brightness, which may be reasonably conceptualized as manifestations of top-down enhancements of sensory gain. If these percepts are themselves driven by prior expectations in the setting of excessive sensory noise, they may be thought to be reflective of perceptual alterations experienced during acute SP dosing. Consistent with this, our two most PPA-predictive SP use patterns are those that result in more intense acute effects.^60,61^ Persistent perceptual abnormalities arising after SP use may be thought of as psychotic-like perceptual phenomena for which there is an identifiable inciting event.

Our study has several limitations worth noting. First, despite the favorable size of our sample relative to other computationally-oriented behavioral studies^26,29,69^ this may be a small sample for detecting behavioral and computational effects associated with a broad range of SP use patterns and personal histories cross-sectionally. This is reflected in our mediation model estimates’ greater uncertainty. Our sample may also be noisy: convenience sampling from online community fora reduces our results’ generalizability and introduces noise due to non-compliance, inattentiveness, confusion, and lack of environmental control, ^70^ although a similar sample performing similar tasks was found to perform in a hardware-invariant manner.^26,29^ This sample may also be slightly atypical, demonstrating a bias towards positive experiences with SPs, higher dose and more frequent SP use, and extensive polydrug use. Most participants also had exposure to other HPPD-associated psychotomimetics^37^ that may have contributed to their PPA profile, although PPA groups did not systematically vary on psychotomimetic exposure. Our retrospective SP use metrics are also likely to feature considerable recall error. Lastly, empiric detection probabilities were significantly higher than expected, suggesting some error in visual threshold calculations. Though we account for this error by using empiric probabilities from an out-of-set sample, this necessarily assumes equal error in thresholding across participants.

In their character and frequency, the PPAs examined here resemble those present in the earliest phases of the psychosis prodrome, characterized by sensory distortions and intensity enhancements rather than fully-formed hallucinations.^71–73^ Whether and how similar processes may be at play in spontaneously-occurring early psychotic phenomena remains unclear, as does the interaction between mechanisms driving early symptom development and those arising from psychedelic or other psychotogenic exposures. It also remains unclear from this cross-sectional sample whether variables related to sensory noise and cognitive biases above constitute risk factors, correlates of active symptomatology, or compensatory effects after SP exposure, and inferring the lasting effects of acute exposure weeks to years afterward is difficult at best. Future work will require assessment of these variables before, during, and after prospective SP exposure.

## Acknowledgments

The authors would like to thank Kayla Morgan, Gabby Hernandez, and Sara Azzi for assisting in data collection, quality control, and advertisement. We thank Dustin Bufford and Mica Chayro for generously offering to advertise our study. We also thank all of the reddit moderators who allowed us to advertise our study: r/psychonaut, r/LSD, r/DMT, r/Mescaline, r/5MeODMT, r/Ayahuasca, r/HPPD, and r/LSD, r/Shrooms, r/Drugs, and r/RationalPsychonaut. We especially thank the consortium of moderators at r/Drugs who created a centralized review system for advertising research studies across multiple subreddits. We also want to thank Bluelight.org and Dr. Monica Barratt for allowing us to advertise on their website and Discord. We want to especially thank the DMT Nexus and Shroomery.org who promoted the study on their websites.

We thank Catalina Mourgues for her analysis advice throughout the study.

We wish to thank Silas Keeter, who assisted in deployment of the VCH task online. Special thanks to Adam Wallis who programmed a task that was not used in this current publication but hastened data collection for this study.

We also want to thank the Shuster and Greenwald families as well as Galen Posch, Jacob Johnston, Ryan Campbell, Katie Stimler, Karlos Mate Piovanetti, and Adam Ismail for testing out the survey and online task.

Special thanks to anonymous participants who helped refine our SP-fraud-detection questions.

Finally, we would like to thank all of our participants who volunteered their time and experiences for this study.

## Author contributions statement

M.G., A.P.—conceived the study.

M.G.—designed the study; produced the manuscript draft and figures, performed survey distribution and data collection, analyzed outcomes, interpreted results, and coordinated authors.

A.P.—acquired funding; designed VCH task and conceived of HGF modeling of VCH data; and was directly involved in all stages of result interpretation; revised and edited manuscript.

E.K. designed, tested, and implemented online VCH task.

M.G., A.P., D.F., S.W.N., P.T.W, S.I, E.K.––revised and edited manuscript.

A.B., P.T.W, D.F., S.W.N, S.I.—gave expert guidance in statistical modeling, diagnostics, and interpretation. All authors substantially contributed to the discussions and manuscript drafting and approved the final version of the manuscript.

## Data Availability Statement

The datasets generated during and/or analysed during the current study are available at this Open Science Framework (OSF) link: https://osf.io/8bsc2/overview?view_only=d3b965144d2449229dac1cdd273a647a under the “datasets” repository.

Code sufficient to reproduce the results are available at this Github repository link: https://github.com/maxsupergreenwald/Computationally-Characterizing-Psychedelics-Persisting-Perceptual-Abnormalities---Public-Code.

The authors welcome contact with questions, feedback, collaboration proposals, etc.

