## Supplementary Materials for "Persisting perceptual abnormalities in psychedelic users are associated with conditioned hallucinations and impaired sensory processing"

**Supplementary Material**

Lifetime SP-associated PPA Screener

Questions were adapted from an online survey study by Baggott and colleagues.^1^ Participants first received the instructions:

*Not counting times when (1) you were under the influence of another psychoactive drug; (2) you were in a trance, falling asleep, waking up, or had not slept in a long time; or (3) you were experiencing known symptoms of a psychotic-spectrum disorder (schizophrenia, bipolar disorder, schizoaffective disorder, schizophreniform disorder, etc.),*

*Have you ever had a period in your life when—FOLLOWING USE OF A SEROTONERGIC PSYCHEDELIC, IN THE DAY(S) AFTER YOU WERE NO LONGER HIGH—you NEWLY experienced any of the following visual effects?*

Before then selecting from the following 12 items:

| *#* | *Visual Effect* | *Yes* | *No* |
| --- | --- | --- | --- |
| *1* | *Halos or auras around things* | *☐* | *☐* |
| *2* | *Stationary things appearing to move, breathe, grow, or shrink* | *☐* | *☐* |
| *3* | *Moving objects appear to not be moving* | *☐* | *☐* |
| *4* | *Afterimages left behind moving objects* | *☐* | *☐* |
| *5* | *Brighter or more intense colors* | *☐* | *☐* |
| *6* | *Seeing patterns or textures that aren't there with eyes open* | *☐* | *☐* |
| *7* | *Seeing patterns or textures with eyes closed ** | *☐* | *☐* |
| *8* | *Seeing objects that aren't really there* | *☐* | *☐* |
| *9* | *Increased intensity in oscillating or flashing lights (TV, light bulbs, etc.)* | *☐* | *☐* |
| *10* | *Distortion, movement, or patterns in grids, gratings, or closely spaced lines* | *☐* | *☐* |
| *11* | *Noticing more things in your surroundings ** | *☐* | *☐* |
| *12* | *Things just looked/seemed different ** | *☐* | *☐* |

** Not included in Baggott et al. 2012.*

Those who endorsed any SP-associated PPAs received follow-ups:

*1. Which effect was the most vivid/intense?*

1. *I have never experienced any of the above visual effects*
2. *Halos or auras around things*
3. *Stationary things appearing to move, breathe, grow, or shrink*
4. *Moving objects appear to not be moving*
5. *Afterimages left behind moving objects*
6. *Brighter or more intense colors*
7. *Seeing patterns or textures that aren't there with eyes open*
8. *Seeing patterns or textures that aren't there with eyes closed*
9. *Seeing objects that aren't really there*
10. *Increased intensity in oscillating or flashing lights (TV, lightbulbs, etc.)*
11. *Distortion, movement, or patterns in grids, gratings, or closely spaced lines*
12. *Noticing more things in your environment*
13. *Things just looked different*

*2. What is the longest you experienced any of these visual effects after a serotonergic psychedelic?*

- *I've never experienced any visual effect the day after taking a psychedelic*
- *< 1 day*
- *1 - 3 days*
- *3 days - 1 week*
- *1 week - 1 month*
- *1 - 6 months*
- *6 months - 1 year*
- *>1 year*

*3. Were these visual effects constant, present for brief spurts (minutes or seconds), or came and went in longer periods (hours to days)?*

- *I have never had visual effects starting the day after taking a psychedelic*
- *Brief spurts (seconds to minutes)*
- *Longer periods (hours to days)*
- *Constant*

*4. About how many times had you taken psychedelics before you first noticed any of these visual effects.*

Cardiff Anomalous Perceptions Scale (CAPS)^2^ Visual Items

Participants were instructed:

*Please do NOT include experiences you've had while under the influence of any psychoactive, spiritual, or ceremonial drug. Please think about your experiences in the past MONTH when answering these questions!*

Before being presented with the following Yes/No questions:

- *Do you ever see shapes, lights or colors even though there is nothing really there?*
- *Do you ever find the appearance of things or people seems to change in a puzzling way, e.g. distorted shapes or sizes or color?*
- *Do you ever look in the mirror and think that your face seems different from usual?*
- *Do you ever have days where lights or colors seem brighter or more intense than usual?*
- *Do you ever think that everyday things look abnormal to you?*
- *Do you ever see things that other people cannot?*

For all endorsed items, participants were asked:

- *How distressing is this for you?*

1. *Not at all distressing*
2. *Slightly distressing*
3. *Somewhat distressing*
4. *Firmly distressing*
5. *Very distressing*

- *How distracting is this for you?*

1. *Not at all distracting*
2. *Slightly distracting*
3. *Somewhat distracting*
4. *Firmly distracting*
5. *Very distracting*

- *How often does this happen?*

1. *Hardly happens at all*
2. *Does not happen often*
3. *Happens sometimes*
4. *Happens regularly*
5. *Happens all the time*

SP use questionnaire

The primary SP-use variables used for analysis were derived from the following questions:

- Lifetime SP uses: *NOT including microdoses(!) -- How many times in your life have you used serotonergic psychedelics?*
- Age of first SP use: *At what age did you first use a serotonergic psychedelic?*
- Average SP dose used: Participants rated the approximate percentage of their lifetime SP uses based on ordinal categories are based on degree of subjective, experiential impairment (as is often used in community forums)^3^ as well as an approximate phenomenology-to-dose guide for LSD ^4^ recommended to the authors by users. The categories were described as:
  - Microdose (20-30ug; barely detectable or "threshold" drug effect)
  - Low dose (40-60ug; detectable but very mild drug effect; hallucinations barely present)
  - Medium dose (90-250ug; full psychedelic effect; many hallucinations)
  - Heavy dose (300-500ug; strong psychedelic effect)
  - Very heavy dose (700ug+; out-of-body experiences and beyond)

And participants were then asked, for each DOSE: *Of all the times you have used psychedelics, what percent of uses would you consider -- based on subjective criteria below -- "DOSE"?* Average SP dose used was then calculated as a weighted average (based on user estimated proportions) of each dose after converting the dose categories into their approximate LSD microgram equivalents shown above.

The text “Serotonergic Psychedelics” was consistently linked to this Wikipedia-curated lis of known psychedelics with significant Serotonin 2A Receptor agonist activity.^5^

Users also reported their last dose using the above ordinal scale for subjective dose and a fully subjective VAS scale from “not detectable” to “strongest imaginable dose.” Users also reported their preferred and most recent SP, the approximate number of uses in the past 6 months, and past month as well as the days since their last use, age of first use, frequency of use during their heaviest period of use and duration of their heaviest use-period.

VCH Task Quality Control Procedures

*VCH data exclusion criteria*

Participants’ VCH task data were excluded if:

1) Detection probability failed to confidently (p < 0.05) increase with contrast intensity as determined by OLS regression—suggesting that the participant either was randomly responding or the threshold determination had failed.

2) There were fewer than three reversals during the QUEST^6^ thresholding procedure—suggesting that a participant had reached an asymptote in the QUEST procedure and hence failed to reach their true visual threshold.

3) There was a set of more than 40 contiguous trials in which the same response was selected *and* *during which* exhibited an implausibly fast reaction time (<450 ms) or significantly (200%) worse hit rate—suggestive of a significant (>10%) portion of the task in which a participant was not engaged.

*Measures of VCH task engagement*

We considered six specific task quality metrics for the purpose of assessing whether decreased decision precision (*β*) estimates were the result of poor task engagement (Supplementary Fig. **S7**):

1. Self-reported, likert-scale ratings of “I did the best I could during the games” and “I was not distracted by my phone or other things during the games” ranging from “strongly disagree” to “strongly agree” on a five-point likert scale.
2. The number of trials where the stimulus had to be re-presented due to non-response the first time, potentially indicating distraction.
3. The error in QUEST-estimated visual threshold and empiric probability of detection in the 75% intensity trials, potentially indicating that participants did not engage properly during the threshold-determination procedure.
4. Two separate composite scores testing for random responding in the form of contiguous trials in which participants selected the same response (“Yes I see it” or “No I don’t see it”). To distinguish such events from authentically identical responses, these composite scores considered the extent to which reaction time or accuracy decreased during this contiguous response sequence. Hence, the final composite scores were calculated as the longest contiguous streak of identical responses (perseveration) multiplied by decrease in median reaction time or accuracy—Z scored. Accuracy was calculated as hit rate + correct reject rate / VCH rate + miss rate.

Fraudulent Response Criteria

To prevent fraudulent responses, participants were not invited to the study if they used a voice over internet protocol (VOIP) phone number, had the same internet protocol (IP) address as two or more existing records, or had a VPN used by records otherwise deemed to be fraudulent via criteria discussed below. After study completion, participants’ responses were screened for evidence of fraud based on inconsistencies or fraudulent responses to “attention check” and “challenge” questions.^7^ Fraudulent responses were flagged, excluded from analysis, and denied payment. Participants were asked to rate their honesty, effort, attention, and degree of distraction at the end without penalty. Participants were removed from data analysis if they endorsed “disagree” or “strongly disagree” to the statement “I answered the questions truthfully” or if they endorsed “disagree” two or more of the following: **“**I paid close attention to the questions,” “I did the best I could on the tasks,” or “I was not distracted by my phone or other things during the surveys and tasks.”

In addition to the screening above, we excluded responses for:

Failed “**Challenge questions**” including:

1. Differences in identical questions between screening and main survey including:
   a. Self-reported race (unless having endorsed “multiracial” on either, in which case participants were coded as “multiracial”).
   b. Self-reported age (difference of up to two years tolerated).
   c. Most recently used SP.
2. Endorsement of any of the following bogus side effects in response to the question “*Which of these common side effects do you USUALLY (>75% of the time) experience while using psychedelics?*”
   a. *Blacking out (falling asleep and losing memory)*
   b. *Small, "pin-point" pupils*
   c. *Hunger and "binge" eating (eating much, much more than you usually would)*
   d. *Severe constipation lasting days after use*
   e. *Gum Bleeding*
3. Endorsing an incredibly unlikely or impossible route of administration for their selected “primary” SP including:
   a. Mescaline: *Smoked, snorted, injected, or lozenge.*
   b. LSD: *Smoked, vaporized, snorted, or injected.*
   c. Psilocybin: *Smoked, vaporized, snorted, injected, or blotter/tab.*
   d. DMT: *Pill (pure DMT), injected, lozenge, or blotter/tab.*
   e. 5-MeO-DMT: *Injected, blotter/tab, pill, or suppository.*

**Inconsistent** answers including:

1. Reporting fewer SP uses within a time window (e.g. the past year) than a more recent time window (e.g. the past month).
2. Reported most recent macrodose being during a time window they reported 0 SP use in (e.g. four months ago when reporting 0 SP uses in the past 6 months) or reporting SP uses during a time period more recent than their most recent use (e.g. last use three months ago, but two uses in the past month).

And finally, failure to select correct response to any of the following five “**attention check**” questions:

1. *Please select "rabbit" to show that you are still paying attention*
2. *I am paying attention right now and reading each entire question so I will select option five.*
3. *Please enter "1" to show you are still paying attention*
4. *Please select "Very Often or Always True" to show you're paying attention* (embedded within a series of likert scale ratings in which *"Very Often or Always True"* was an option)
5. *Please select “Yes” to show you’re paying attention*

For all of the above cases, participants were allowed to schedule a Zoom call and explain/modify their

responses and could have their modified data included at researchers’ discretion.

See **Supplementary Table S4** for the breakdown of fraud indicators in our sample.

Hierarchical Gaussian Filter (HGF) Model: Bayesian Workflow for Computational Psychiatry

To assess face-validity and identifiability of models, we performed a prior-based parameter recovery. For all four models described above (two and three-level with and without empiric likelihoods), we randomly sampled 500 parameter values from our prior distributions, generated synthetic behavioral data, and then inverted each model using markov-chain monte-carlo (MCMC; see next section “Additional details” below) and calculated Bayesian Information Criterion (BIC) values from the maximum likelihood over the posterior. Parameter recovery was assessed by plotting generative parameters against median estimates for each model (**Supplementary Fig. S3)**, and model identifiability was assessed based on the proportion of samples (out of 500 generative samples) in which the model used to generate the data had the lowest BIC after model inversion (Supplementary Fig. **S3)**. Across all models, *ν* and *β*‘s recovery were excellent (*r* > 0.8; **Supplementary Fig. S3**), while *ω*’s was acceptable (*r* ~ 0.4), and *ω_3_*’s was unrecoverable (*r* < 0.1; all posterior estimates within 0.01 of the prior; **Supplementary Fig. S3**).

The two-level HGF with empiric detection likelihoods was selected based on having the greatest expected posterior frequency (Ef; 0.805) and protected exceedance probability (PXP; 1.00) in Random-effects Bayesian model selection (RFX-BMS) described in detail elsewhere (**Supplementary Fig. S3**).^8,9^ This approach is appropriate for comparing the two versus three-level models. However, it may not be the definitive test for adjudicating between the empiric versus QUEST-estimated detection probabilities because model comparison presumes identical observations being explained, and detection probabilities are arguably seen as HGF inputs. We ultimately decided to proceed with the empiric detection probabilities for the following reasons: 1) deviations from QUEST estimated detection probabilities were large enough to be behaviorally relevant (the minimum difference between the QUEST-estimate and lower bound of the 94% confidence interval of empiric detection in the target-present conditions was >13%; **Supplementary Fig. S2**). 2) posterior predictive checks using the nominal, QUEST-derived detection likelihoods significantly deviated from actual detection behavior (**Supplementary Fig. S2**), 3) empiric detection probabilities were not significantly different between this sample and the two subsets of a separate sample, suggesting consistent error in QUEST-estimation (**Supplementary Fig. S2**), 4) detection probabilities did not significantly change over the course of the experiment, suggesting that a grand mean was appropriate as the “ground truth” detection probability, and finally 5) prior work in voice-hearing populations also used empiric detection probabilities for similar reasons–because stimulus intensity itself was not measurable outside of a carefully-controlled laboratory experiment.^10^ Still, in acknowledgement of the uncertainty regarding this modeling choice, we include the primary regression and mediation results for *β* and *ν* estimates derived from the HGF model using nominal, QUEST-derived detection probabilities as likelihoods (**Supplementary Table S5**). With these model estimates, lower *β* was still associated with SP-related PPA history. and *ν* still was not. However current PPAs were not associated with *β*, and instead were much more strongly associated with *ν.* However, *ν* still did not mediate average SP dose effects on current PPAs, and *β* was plausibly influenced by average SP dose. Taken alongside the compelling reasons for using the empiric detection probabilities, these results should be considered as evidence for *some* role for prior signaling in SP-related PPAs, as discussed in the **Discussion**. We remain confident in our ultimate interpretation being influenced primarily by model results from HGF models using the empiric likelihoods.

Once we had posterior estimates from the 2-level HGF, we employed posterior-based parameter recovery and qualitative posterior predictive checks (PPCs)^11^ as final model validation. In parameter recovery, we took the median parameter estimates for each participant and generated 10 synthetic behavioral datasets and then refit the model before then plotting generative vs. recovered parameters. We verified excellent recovery rates (*r* = 0.936, 0.832, 0.764 for *ν*, *β,* and *ω*, respectively; **Supplementary Figure S3**). We also verified parameter identifiability via correlations between generative parameters and *other* recovered parameters, where the highest correlation was between generative *ν* and recovered *ω* (*r* = 0.43). Correlations between generative *ν* and *β* were much smaller (*r* = -0.13; **Supplementary Figure S3**). For PPCs, we simulated one dataset per participant based on the median estimates for each of their three HGF parameters, and then evaluated model validity based on: 1) increasing detection probabilities with stimulus strength and 2) decaying detection probability over the course of the experiment. Both these criteria were well-met; overall, HGF simulated data tended to over-predict VCH rate and under-predict hit rate (Supplementary Fig. **S3**).

Hierarchical Gaussian Filter (HGF) Model: Additional details

*Generative Model*

The generative model used by the HGF describes the dynamics of the set of continuous, categorical or binary states in the environment that are tracked by the HGF. Continuous states evolve as Gaussian random walks, a flexible assumption that is applicable without specific knowledge of the environment’s true dynamics. The parameters of these Gaussian random walks (most importantly the volatility ω) are influenced by other states- estimating those states with Bayesian inference therefore allow for dynamically estimating volatilities, making the HGF particularly suited for doing inference on volatile environments. Binary and categorical states evolve according to Bernoulli and categorical probability distributions, respectively; since the probability parameters of these distributions are controlled by higher-level estimated continuous states, the HGF can also learn the probabilities of discrete outcomes.

The HGF is now implemented as a network of nodes, with each node representing one of the states tracked by the HGF. In the specific generative model structure used in this experiment, a binary observation u is a noiseless reflection of a binary state *X1* which encodes whether a signal was present. *X1* is controlled entirely by its value parent, a continuous state *X2* which encodes the probability of *X1* being 0 or 1. *X2* itself has a volatility parent *X3* which encodes the changing volatility of *X2*. Since they evolve as Gaussian random walks, *X2* and *X3* additionally depend on their own previous states. This dependency is governed by their respective constant volatilities *ω* and *ω_3_*. Other parameters exist- continuous nodes can have constant drifts *ρ* as well as an autoconnection strength λ that determines the influence of its previous state, and coupling strengths *κ* can be modulated to control how much various nodes affect each other- but they are not used (i.e. fixed to default values) in the model for this experiment. HGF models with this type of generative model are known as variations on the classic binary HGF. Notably, the volatility parent *X3* can be removed from the generative model, so that no phasic volatility is estimated. Models without and including *X3* are then known as 2-level and 3-level binary HGF models. Both types of HGFs are used and compared here, but after model comparison, the 2-level version is used for the main results

*The Inference Process of the HGF*

The HGF’s generative model describes its assumptions about the dynamics in the environment, and how it generates observations. This generative model is then inverted based on received observations, in order to form beliefs *ϑ* about each environmental state *x* in the generative model on an observation-by-observation basis. This - the inference process - is accomplished by using variational Bayesian inference, where the Bayesian posterior belief is approximated by minimizing a variational free energy over possible approximate posteriors. Due to the specific composition of its generative model, the HGF can per-form variational inference quickly using a set of single-step update equations that consist of belief updates based in precision-weighted prediction error signals. Here, Gaussian predictions (with mean *µ*ˆ and precision *π*ˆ) are formed and passed through the network of belief states, from *x*_3_ to *u*. For each node, and in the opposite order, a precision-weighted prediction error *ε* is calculated, and used to update the Gaussian posterior (with mean *µ* and precision *π*) of its parent. This single-step update is made possible by a quadratic approximation to the free energy landscape - there are multiple methods for performing this approximation, but we here used the original algorithm.^12^

Ordinarily, HGF models receive noisy but definite observations from the environment, based on which they learn its probabilistic contingencies. In the conditioned hallucinations task, however, the learning is based on a subjectively experienced percept, which requires the HGF to be augmented with a model of the process of forming percepts. Here, a perceptual posterior belief is formed by combining the stimulus intensity *s* (i.e. likelihood) with the expectation *µ*ˆ (i.e. prior) provided by the HGF, where the balance between the two is controlled by a weighting parameter *ν*. This belief then results in a subjective percept *u*, with a precision parameter *β* controlling the stochasticity of this process. *u* is then passed to the HGF, which in turn updates beliefs and predictions about future observations.

Notably, instead of relying solely on the experimentally imposed conditions for stimulus probabilities, we incorporated empirical response rates derived from participants’ behavior using an out-of-sample dataset. This allowed the model to better reflect participants’ actual learning trajectories and belief updates, accounting for potential deviations from the experimental contingencies. Thus, the HGF model used in this experiment in total has four free parameters Θ which are estimated:

*ω*_2_ *∈* [*−∞, ∞*]: the expected (log) tonic volatility of the probability. This parameter controls the baseline uncertainty of probability beliefs and in turn their learning rate so that higher *ω*_2_ values lead to faster updating of the estimated outcome probability.

*ω*_3_ *∈* [*−∞, ∞*]: the expected (log) tonic volatility of the phasic volatility. This parameter controls the baseline uncertainty of volatility beliefs and in turn their learning rate so that higher *ω*_3_ values lead to faster updating of the estimated phasic volatility.

*ν ∈* [0*,* 1]: the relative weighting of the prior against the likelihood when forming perceptual beliefs. Higher *ν* values lead to stronger weighting of the prior (i.e. the prediction supplied by the HGF) relative to the likelihood (i.e. the stimulus intensity).

*β ∈* [0; *∞*]: the consistency of perceptual decisions with evidence. Higher *β* values leading to percepts that are in accordance with perceptual beliefs, independent of both prior and likelihood.

In addition, the model produces a set of inferred trial-by-trial belief states *ϑ*. Of specific relevance for the analysis in this article are:

*b ∈* [0*,* 1]: the posterior perceptual belief, after combining stimulus and prior expectations.

*µ*_2_[*−∞, ∞*]: the (logit) posterior belief about the outcome probability, with a higher value meaning a higher expected probability of a signal, and with 0 corresponding to a probability of 0.5.

*ε*_2_ *∈* [*−∞, ∞*]: the precision-weighted prediction error. A value of 0 indicates a perfect correspondence between prediction and observation, higher values means that the observation was above the prediction and vice versa.

*HGF Parameter Estimation*

MCMC for Bayesian inference of individual parameters was performed using the No U-Turn Sampler (NUTS)^13^ for 1000 iterations over 4 chains. We confirmed R̂ values within 1 ± 0.01 between all chains for any estimate incorporated into any primary or supplementary analyses.

For parameter priors, we used posterior parameter distributions from a prior HGF designed for CH data^10^ that were slightly more uncertain and truncated in regions that are undefined for the HGF:

*β ∼ N (3.41, 1) for β > 0.001,*

*ν ∼ N (0.7265, 1) for 1 ≥ ν ≥ 0,*

*ω_2_ ∼ N (−5.1683, 1) for ω ≤ −0.5.*

*ω_3_ ∼ N (−6, 1) for ω ≤ −0.5.*

Here, *N* (*µ, σ*^2^) denotes a normal distribution with mean *µ* and variance *σ*^2^, truncated to the specified region for each parameter.

*Perceptual response model*

We use the unit square sigmoid function to transform the belief *b_t_* at trial *t* into a probability *P* (*u_t_*) of making a subjective sensory observation *u_t_* of either 0 or 1 (signal absent or present). The action precision parameter *β* controls the precision of this probability, with higher values leading to probabilities further from 0.5 and closer to the extremes:

(S1) $P(u_{t})= \frac{{b_{t}}^{\beta}}{{b_{t}}^{\beta}+ (1 - b_{t})^{\beta}}$

The belief *b_t_* is calculated as a weighted combination of the prior (i.e., the prediction about the observation *µ*ˆ_1_*_,t_*, which is provided by the HGF before receiving a stimulus), and the strength of the signal *s_t_* (i.e., the likelihood), with the relative weighting between the two controlled by the prior weighting parameter *ν*:

(S2) $b_{t}= \hat{\mu}_{t,1}+\frac{1}{1+\nu}\cdot(s_{t}- \hat{\mu}_{1,t})$

Bayesian regression model specification and inference approach:

For models examining predictors of SP-associated PPA history, we used a Bernoulli likelihood with logit link. For those examining predictors of current CAPS visual item count—where the non-endorsement rate is unusually high—we used a hurdle negative binomial distribution with a log link for the mean and a logit link for the hurdle. Hurdle models are effectively mixture models that involve both a Bernoulli likelihood for whether or not a variable is zero (the “hu” or “hurdle” component) as well as the primary model estimating change in mean count for those with nonzero values (negative binomial likelihood in our case).^14^ For both response variables, we infer association based on the probability that increasing the predictor by one standard deviation causes a consistent, nonzero change in the response variable.^15,16^ To do this, we simulate 1000 predicted values from the posterior predictive distribution for each of 16,000 joint posterior parameter draws (the mean and *hu* coefficients), at the mean and mean + 1 SD predictor values, and then taking the proportion of draws in which the mean predicted value is higher at the larger predictor values. We report the proportion of simulated differences that are positive or negative as a probability of effect—the equivalent of a p-value.^17^ We also report 94% Highest Density Interval (HDI) values of this difference. This approach allows us to incorporate predictors’ effects on multiple parameters (e.g. the mean and “hu”) while maintaining easily interpretable estimates. Raw beta coefficients and probabilities of effects are summarized in **Supplementary Table S6 and S7**.

Bayesian Regression Model Priors

All slopes were fitted with flat priors. The following are the default priors for BRMS v. 2.23.0:

Hurdle negative binomial (CAPS Vision):

- Intercept: β_0_ ∼ Student-*t*(3, *μ*, 2.5)
- Hurdle submodel intercept: α_0_ ∼ Logistic(0,1)
- Shape: ϕ ∼ Inv-Gamma(0.4, 0.3),ϕ > 0

Zero-inflated beta (VCH Rate):

- Intercept: β_0_ ∼ Student-*t*(3, *μ*, 2.5)
- Zero submodel intercept: α_0_∼Logistic(0,1)
- Phi: ϕ ∼ Gamma(0.01, 0.01),ϕ > 0

Bernoulli (PPA History):

- Intercept: β_0_ ∼ Student-*t*(3, *μ*, 2.5)

Gamma (*ν*):

- Intercept: β_0_ ∼ Student-*t*(3, *μ*, 2.5)
- Shape: ϕ ∼ Gamma(0.01, 0.01), ϕ > 0

Model Diagnostics

We confirmed minimal (i.e., p > 0.05) violations of uniformity (via Kolmogorov–Smirnov test), dispersion, or homoskedasticity (via quantile regressions against scaled residuals). Finally, we inspected Q-Q plots of scaled residuals against the uniform distribution as well as plots of scaled residuals against SP, VCH, and HGF predictors for unacceptable residual patterns.

Model diagnostics for CAPS vision and PPA history univariate models are available at this OSF link

“https://osf.io/zgv8p/overview?view_only=281abd88672f45f4939852d79bc04c58”

under “single_path_caps_vision” and “single_path_hppd_binary,” respectively. Each page comprises a single predictor’s effect and features: (**a**) posterior predictive check plots—assessed by comparing the dark line (the actual data’s distribution) to light blue lines (the posterior-simulated distributions) for similarity in location, spread, skewness, and tails; (**b**) trace plots of Markov Chains for all coefficients assigned to the predictor, assessed for a “fuzzy caterpillar” appearance with no obvious trends and heavy overlap; (**c**) the primary figure returned by the simulateResiduals() function in DHARMa,^18^ featuring (**left**) a QQ-plot assessed for detecting deviations from the specified likelihood (a perfect fit would be the data matching the straight line) along with printed results of Kolmogorov–Smirnov test of uniformity (detecting the same), over/underdispersion, and outliers, and (**right**) a quantile regression rank transformed model predictions against the scaled residuals—assessed by the degree to which the three regressions are flat and centered at the expected percentile; and (**d**) another quantile regression (where an ideal model results in three flat lines) of scaled residuals plotted against the primary predictor to assess for heteroskedasticity, linearity assumption, and/or other evidence of misspecification.

Model diagnostics for average SP dose → CAPS vision and age of first SP use → PPA history mediation models are available at this OSF link:

“https://osf.io/7se4k/overview?view_only=5b24779e998342f485ece3d279e946b4”

under “mediation_models_caps_vision” and “mediation_models_hppd_binary,” respectively. Each page comprises a single model and features: (**top**) posterior predictive check plots for (**left**) the response variable (CAPS vision or PPA history) and (**right**) the mediator (VCH rate, detection threshold, or decision precision parameter); (**2nd row**) trace plots of Markov Chains for all coefficients assigned to the predictor; (**third row**) the primary figure returned by the simulateResiduals() function in DHARMa for (**left**) the SP variable + mediator → response variable model and (**right**) the SP variable → mediator model; and (**bottom**) quantile regressions of the DHARMa residuals from (**left two**) the response variable model and (**right**) the SP variable → mediator model plotted against the SP variable and (for the response variable model only) the mediator.

The DHARMa^19^ package provides a means of assessing model specification/assumption violation for generalized linear models as typically handled by residuals plots with normal linear models. Interested readers are referred to the CRAN page (https://cran.r-project.org/web/packages/DHARMa), but, briefly, DHARMa calculates a “scaled residual” for each actual data point as the quantile of that observation within the posterior-simulated distribution defined by our model—a 0 to 1 value representing the proportion of the simulated dataset that is smaller than the observation.

We note that models predicting CAPS vision endorsements—particularly mediation models—tended to show varying signs of an elevated 75th quantile regression line compared to model predictions, suggesting that models consistently under-predicted CAPS scores for those with the most (>3) symptoms in a manner that did not vary with any of our predictors. While this evidence of misspecification should caution interpretation, it does not, in and of itself, suggest specific bias requiring modification of our results. Importantly, other models that did not raise these misspecification flags still exhibited the same effects (Supplementary Fig. **S4**), making us comfortable that our inferences were reasonable. The most concerning finding here was that the average SP dose → VCH rate → CAPS vision had this issue in every model except the univariate model, where the estimated probability of mediation was lower (~89%), *potentially* suggesting that the primary results may have overestimated confidence, though not the primary inference, of this effect.

For PPA history, we note an elevated 75th quantile regression line for VCH rate and a depressed 25th quantile regression line for hit rate in the 75% threshold trials in the univariate models. However, as above, multiple other models that did not have these misspecification concerns but still had the same effects, (Supplementary Fig. **S4**) leading us to conclude that our inferences were not biased. In mediation models of age of first SP use effects on PPA history, we note that the VCH rate mediator model predictions regressed towards the mean; underestimating for high probabilities and overestimating for low ones. This did not appear to be related to any particular bias; there was again an elevated 75th quantile regression line for VCH rate. This same pattern was evident for *ν*.

**Supplementary Tables**

| **Ineligibility Reason** | **Total** |
| --- | --- |
| Fraud-associated phone # or IP address | 545 (72.0%) |
| No computer | 21 (2.8%) |
| Non-English speaking | 1 (0.1%) |
| >65 years old | 1 (0.1%) |
| <18 years old | 2 (0.3%) |
| Neurocognitive Impairment | 18 (2.4%) |
| Epilepsy | 2 (0.3%) |
| Active intoxication | 46 (6.1%) |
| 0 RAVEN score | 10 (1.3%) |
| Recent heavy cannabis use | 96 (12.7%) |
| Atypical psychedelic more recent than SP | 43 (5.7%) |
| No SP Use | 5 (0.7%) |

**Supplementary Table S1: Summary of reasons that screened records were found ineligible.** Phone numbers and IP addresses were considered fraudulent based on: 1) sharing an IP organization or location with high rates of participants who failed QC (see **Supplementary Table S2** and “Fraudulent Response Criteria” in **Supplementary materials**), 2) using a voice-over-internet protocol through the Sinch corporation, and 3) Fraud scores and “recent abuse” designations from IPQualityScore.com. A score of 0 on the RAVEN was taken to indicate non-engagement in screening. See **Methods** for our determination of “recent heavy cannabis use”. Atypical psychedelics discussed in screening included: dextromethorphan, ketamine, phencyclidine, 3,4-Methylenedioxymethamphetamine, salvinorin A, scopolamine, and ibogaine.

| **Variable** | **ρ** | **p** |
| --- | --- | --- |
| ***Demographic Characteristics*** | | |
| Age | -0.235 | .007 |
| Sex: Male | -0.009 | .921 |
| Sex: Female | +0.009 | .921 |
| Race: White | +0.119 | .179 |
| Race: Latino/a | -0.101 | .251 |
| Race: Asian | +0.035 | .690 |
| Race: Multiracial | -0.159 | .072 |
| Race: Am. Indian/Alaska Native | +0.077 | .381 |
| Race: Unknown/Prefer not to say | +0.077 | .381 |
| Location: United States | -0.237 | .007 |
| Location: Europe | +0.277 | .001 |
| Location: Oceania | +0.009 | .921 |
| Location: Latin America | -0.121 | .171 |
| Location: Other | +0.074 | .402 |
| Education (9-level) | -0.176 | .046 |
| Education (3-level) | -0.118 | .182 |
| Education (6-level) | -0.167 | .057 |
| Raven's Progressive Matrices (9-item) | +0.014 | .875 |
| ***Psychiatric History*** | | |
| Any Diagnosis | +0.028 | .755 |
| Psychotic Spectrum | +0.123 | .162 |
| Schizophrenia Spectrum | +0.123 | .162 |
| Unipolar Mood Disorder | -0.028 | .749 |
| Bipolar Disorder (no psychosis) | +0.112 | .205 |
| Substance Use Disorder | +0.205 | .020 |
| Anxiety Disorder | -0.033 | .710 |
| OCD | +0.035 | .689 |
| Trauma-Related Disorder (PTSD/c-PTSD) | +0.036 | .683 |
| Eating Disorder | +0.115 | .193 |
| Personality Disorder | -0.020 | .818 |
| Autism Spectrum Disorder | -0.007 | .939 |
| ADHD | -0.080 | .368 |
| Sleep Disorder | -0.002 | .980 |
| ***Psychiatric Medications*** | | |
| Any Medication | -0.140 | .111 |
| Antipsychotic | -0.121 | .171 |
| Antidepressant | -0.046 | .603 |
| Stimulant Medication | -0.149 | .091 |
| Benzodiazepine | -0.032 | .718 |
| Anxiolytic (Non-Benzodiazepine) | -0.118 | .183 |
| Sedative (Non-Benzodiazepine) | -0.148 | .092 |
| Opioid Antagonist | +0.151 | .087 |
| ***Other Substance Use (Past Month)*** | | |
| Any Substance | -0.030 | .737 |
| Alcohol | +0.017 | .851 |
| Sedative-Hypnotic | -0.075 | .393 |
| Opioid | -0.032 | .718 |
| Cannabis (binary) | -0.028 | .755 |
| Atypical Psychedelics (binary) | +0.095 | .280 |
| Stimulants (binary) | +0.135 | .125 |
| Cannabis (count, past 6 months) | -0.101 | .252 |
| Atypical Psychedelics (count, past 6 months) | +0.159 | .071 |
| Stimulants (count, past 6 months) | -0.055 | .534 |
| Sedative-Hypnotic (count, past 6 months) | +0.012 | .891 |
| ***Other Substance Use (Lifetime)*** | | |
| Alcohol | -0.069 | .433 |
| Sedative-Hypnotic | -0.061 | .491 |
| Opioid | +0.024 | .788 |
| Cannabis | -0.045 | .608 |
| Atypical Psychedelics | +0.039 | .657 |
| Stimulants | +0.064 | .472 |

**Supplementary Table S2: Spearman correlations of demographic/clinical variables with CAPS visual item endorsements for identification of additional covariates for sensitivity analyses.** Note that “substance use disorder” was excluded as a covariate due to low sample size (n = 9), and the 6-level Education variable (which collapsed the lowest and highest education levels with their nearest level → “High school diploma/GED or less” and “Master’s degree or other post-graduate degree”).

| **Predictor** | **PPA(−) Mdn [Q1, Q3]** | **PPA(+) Mdn [Q1, Q3]** | **U** | ***r_rb_*** | **p** | **N** |
| --- | --- | --- | --- | --- | --- | --- |
| ***SP Use Patterns*** | | | | | | |
| Age at First Use (Years) | 22 [18.5, 31] | 19 [17, 23] | 4624.5 | +0.284 | 0.002 | 186 |
| Lifetime SP Uses (Count) | 15 [5, 29] | 20 [6, 43.5] | 3040 | -0.156 | 0.093 | 186 |
| Avg. Dose (LSD μg eq) | 175 [102, 333.125] | 212.5 [145, 324.75] | 3238 | -0.101 | 0.277 | 186 |
| ***VCH Task Behavior*** | | | | | | |
| 75% Threshold | -0.531 [-0.615, -0.458] | -0.574 [-0.641, -0.513] | 3206 | +0.223 | 0.028 | 160 |
| Hit Rate (75% Contrast Trials) | 0.924 [0.875, 0.958] | 0.917 [0.861, 0.958] | 2596.5 | -0.01 | 0.925 | 160 |
| VCH Rate | 0.042 [0.016, 0.075] | 0.059 [0.035, 0.118] | 1948 | -0.257 | 0.011 | 160 |
| ***HGF Estimates*** | | | | | | |
| Prior Weighting (ν) | 0.369 [0.182, 0.675] | 0.392 [0.127, 0.829] | 2607 | -0.006 | 0.956 | 160 |
| Decision Precision (β) | 4.125 [3.612, 4.461] | 3.773 [3.332, 4.183] | 3226 | +0.23 | 0.023 | 160 |
| Contingency Belief Evolution Rate (ω) | -4.661 [-5.113, -3.746] | -4.898 [-5.232, -3.73] | 2793 | +0.065 | 0.520 | 160 |

**Supplementary Table S3: Table of statistics from Mann-Whitney-U tests comparing participants with and without a history of SP-associated PPAs.** The first two columns report the medians followed by the interquartile range for individuals without and with a history of SP-related PPAs. Effect size is captured by the *r_rb_* or rank biserial correlation (interpretable like a correlation coefficient).

| **QC Failure Reason** | **Total** |
| --- | --- |
| Inconsistent Answers | 49 (59.0%) |
| Failed Attention Checks | 12 (14.5%) |
| Failed Challenge Questions | 34 (41.0%) |
| Fraud-associated Phone / IP | 10 (12.0%) |
| **Supplementary Table S4: Summary of reasons for quality-control (QC) failure after study completion. “**Fraudulent Response Criteria” in **Supplementary materials** for further description. Criteria for deeming a phone number or IP address fraudulent are described in **Supplementary Table S1**; participants listed here finished the study before their phone numbers/IP addresses were associated with fraud and had no other major concerns with their record. | |

**a**

| **Predictor** | **β** | **β 94% HDI** | **P(β≠0)** | **Δ** | **Δ 94% HDI** | **P(Δ≠0)** | **N** |
| --- | --- | --- | --- | --- | --- | --- | --- |
| **Prior Weighting (ν)** | **+0.163** | **[-0.497, +0.829]** | **0.673** | **+0.015** | **[-0.052, +0.074]** | **0.673** | **160** |
| **Decision Precision (β)** | **-0.302** | **[-0.654, +0.030]** | **0.951** | **-0.015** | **[-0.023, -0.001]** | **0.951** | **160** |

**b**

| **Predictor** | **β** | **β 94% HDI** | **P(β≠0)** | **hu β** | **hu β 94% HDI** | **P(hu_β≠0)** | **Δ** | **Δ 94% HDI** | **P(Δ≠0)** | **N** |
| --- | --- | --- | --- | --- | --- | --- | --- | --- | --- | --- |
| **Prior Weighting (ν)** | **+0.168** | **[-0.347, +0.632]** | **0.759** | **-1.356** | **[-2.260, -0.441]** | **0.998** | **+0.370** | **[+0.024, +0.751]** | **0.984** | **113** |
| **Decision Precision (β)** | **-0.145** | **[-0.405, +0.109]** | **0.873** | **+0.015** | **[-0.399, +0.424]** | **0.526** | **-0.098** | **[-0.617, +0.090]** | **0.780** | **113** |

**c**

| **Path** | **β** | **β 94% HDI** | **P(β≠0)** | **Δ** | **Δ 94% HDI** | **P(Δ≠0)** |
| --- | --- | --- | --- | --- | --- | --- |
| ***VCH Decision Noise (β)*** | | | | | | |
| **a** | **-0.044** | **[-0.340, +0.268]** | **0.609** | **-0.022** | **[-0.170, +0.134]** | **0.609** |
| **b** | **-0.337** | **[-0.698, +0.003]** | **0.965** | **-0.014** | **[-0.021, -0.003]** | **0.965** |
| **c′** | **-1.218** | **[-2.027, -0.437]** | **0.999** | **-0.066** | **[-0.151, -0.009]** | **0.999** |
| **NIE** | **—** | **—** | **—** | **+0.001** | **[-0.010, +0.013]** | **0.594** |
| **NDE** | **—** | **—** | **—** | **-0.126** | **[-0.209, -0.038]** | **0.999** |
| **Total** | **—** | **—** | **—** | **-0.124** | **[-0.210, -0.037]** | **0.998** |
| **PMed** | **—** | **—** | **—** | **-0.008** | **[-0.145, +0.096]** | **0.593** |
| ***VCH Top-down Bias (ν)*** | | | | | | |
| **a** | **+0.188** | **[-0.173, +0.554]** | **0.836** | **+0.063** | **[-0.061, +0.200]** | **0.836** |
| **b** | **+0.244** | **[-0.425, +0.954]** | **0.746** | **+0.023** | **[-0.041, +0.082]** | **0.746** |
| **c′** | **-1.198** | **[-1.990, -0.413]** | **>.999** | **-0.127** | **[-0.214, -0.041]** | **>.999** |
| **NIE** | **—** | **—** | **—** | **+0.001** | **[-0.007, +0.015]** | **0.658** |
| **NDE** | **—** | **—** | **—** | **-0.126** | **[-0.212, -0.041]** | **>.999** |
| **Total** | **—** | **—** | **—** | **-0.123** | **[-0.209, -0.039]** | **0.999** |
| **PMed** | **—** | **—** | **—** | **-0.012** | **[-0.150, +0.085]** | **0.658** |

**d**

| **Path** | **β** | **β 94% HDI** | **P(β≠0)** | **Δ** | **Δ 94% HDI** | **P(Δ≠0)** |
| --- | --- | --- | --- | --- | --- | --- |
| ***VCH Decision Noise (β)*** | | | | | | |
| **a** | **-0.260** | **[-0.576, +0.066]** | **0.939** | **-0.130** | **[-0.288, +0.033]** | **0.939** |
| **b** | **-0.119** | **[-0.382, +0.149]** | **0.811** | **-0.020** | **[-0.364, +0.081]** | **0.591** |
| **b (hu)** | **-0.115** | **[-0.549, +0.306]** | **0.695** | **—** | **—** | **—** |
| **c′** | **+0.341** | **[-0.229, +0.925]** | **0.885** | **+0.677** | **[+0.074, +1.686]** | **0.997** |
| **c′ (hu)** | **-1.795** | **[-2.841, -0.786]** | **>.999** | **—** | **—** | **—** |
| **NIE** | **—** | **—** | **—** | **+0.004** | **[-0.044, +0.075]** | **0.606** |
| **NDE** | **—** | **—** | **—** | **+0.523** | **[+0.133, +0.953]** | **0.990** |
| **Total** | **—** | **—** | **—** | **+0.537** | **[+0.113, +0.980]** | **0.985** |
| **PMed** | **—** | **—** | **—** | **+0.009** | **[-0.116, +0.170]** | **0.617** |
| ***VCH Top-down Bias (ν)*** | | | | | | |
| **a** | **+0.140** | **[-0.257, +0.558]** | **0.751** | **+0.043** | **[-0.085, +0.178]** | **0.751** |
| **b** | **+0.164** | **[-0.343, +0.674]** | **0.749** | **+0.323** | **[+0.019, +0.666]** | **0.986** |
| **b (hu)** | **-1.455** | **[-2.407, -0.508]** | **>.999** | **—** | **—** | **—** |
| **c′** | **+0.379** | **[-0.212, +0.984]** | **0.909** | **+0.518** | **[+0.148, +0.889]** | **0.998** |
| **c′ (hu)** | **-1.802** | **[-2.926, -0.842]** | **>.999** | **—** | **—** | **—** |
| **NIE** | **—** | **—** | **—** | **+0.018** | **[-0.072, +0.166]** | **0.734** |
| **NDE** | **—** | **—** | **—** | **+0.504** | **[+0.122, +0.877]** | **0.995** |
| **Total** | **—** | **—** | **—** | **+0.538** | **[+0.125, +0.952]** | **0.996** |
| **PMed** | **—** | **—** | **—** | **+0.036** | **[-0.162, +0.344]** | **0.735** |

**Supplementary Table S5: Regression table of univariate Bayesian regression models and multivariate Bayesian mediation models for *β* and *ν* variables estimated from the two-level HGF model using nominal, QUEST-derived detection probabilities as likelihoods**. Beta coefficients (first columns) predict (**a**) mean probability of SP-associated PPA history and (**b**) mean number of visual CAPS items endorsed in the past month to the expected change (Δ) in both variables from a 1 SD increase in the predictor (right half of table). For (**b**) only, center columns also summarize the beta coefficient for the hurdle ("hu") component determining the probability of there being zero CAPS symptoms—negative values indicate a reduced probability of zero symptoms. For the relationship between (**c**) age of first SP use and history of SP-associated PPAs and (**d**) average SP dose and past-month CAPS visual item endorsements, summary statistics of each path in a standard mediation diagram are listed beneath each potential mediating variable explored. The paths are as follows; a: SP use → *β/ν* , b: *β/ν* → PPA variable, c': SP use → PPA variable while controlling for the *β/ν*, NIE: the "natural indirect" effect or "mediated" effect of SP use → mediator → PPA variable, NDE: the combined "natural direct effect" of both c' and (for CAPS vision models only) the hurdle "hu" c', "Total": the combined NIE and NDE effects of SP use → PPA variable, "PMed": the proportion of the Total effect attributable to the NIE. Paths that include “hu” capture the "hurdle" component. Δ represents the expected change in a PPA variable from a 1 SD increase in the predictor or mediator. For CAPS vision models, Δ estimates account for both the β and the hurdle components. The probability of a β coefficient or Δ being non-zero is the primary value used for statistical inference; it is the equivalent to a p-value. HDI = "Highest density interval"—the equivalent of a 94% confidence interval around the mean estimate. See "Bayesian regression model specification and inference approach" in **Supplementary Materials** for more details.

**a**

| **Predictor** | **β** | **β 94% HDI** | **P(β≠0)** | **Δ** | **Δ 94% HDI** | **P(Δ≠0)** | **N** |
| --- | --- | --- | --- | --- | --- | --- | --- |
| ***SP Use Patterns*** | | | | | | | |
| Age at First Use (Years) | -1.147 | [-1.883, -0.399] | **0.999** | -0.120 | [-0.203, -0.042] | **0.999** | 186 |
| Lifetime SP Uses (Count) | +1.167 | [+0.067, +2.420] | **0.987** | +0.095 | [+0.013, +0.178] | **0.987** | 186 |
| Avg. Dose (LSD μg eq.) | +0.108 | [-0.559, +0.794] | 0.609 | +0.010 | [-0.055, +0.077] | 0.609 | 186 |
| ***VCH Task Behavior*** | | | | | | | |
| 75% Threshold | -0.990 | [-1.725, -0.220] | **0.995** | -0.103 | [-0.186, -0.023] | **0.995** | 160 |
| Hit Rate (75% Contrast Trials) | -0.171 | [-0.851, +0.532] | 0.673 | -0.016 | [-0.087, +0.051] | 0.672 | 160 |
| VCH Rate | +0.986 | [+0.072, +1.915] | **0.986** | +0.083 | [+0.014, +0.150] | **0.986** | 160 |
| ***HGF Estimates*** | | | | | | | |
| Prior Weighting (ν) | +0.177 | [-0.485, +0.838] | 0.687 | +0.016 | [-0.047, +0.077] | 0.687 | 160 |
| Decision Precision (β) | -0.749 | [-1.445, -0.075] | **0.981** | -0.077 | [-0.156, -0.009] | **0.981** | 160 |
| Contingency Belief Evolution Rate (ω) | -0.089 | [-0.781, +0.593] | 0.601 | -0.009 | [-0.081, +0.054] | 0.600 | 160 |

**b**

| **Predictor** | **β** | **β 94% HDI** | **P(β≠0)** | **hu β** | **hu β 94% HDI** | **P(hu_β≠0)** | **Δ** | **Δ 94% HDI** | **P(Δ≠0)** | **N** |
| --- | --- | --- | --- | --- | --- | --- | --- | --- | --- | --- |
| ***SP Use Patterns*** | | | | | | | | | | |
| Age at First Use (Years) | +0.019 | [-0.683, +0.747] | 0.521 | +0.800 | [-0.011, +1.674] | **0.967** | -0.146 | [-0.458, +0.176] | 0.811 | 130 |
| Lifetime SP Uses (Count) | +0.233 | [-0.086, +0.554] | **0.932** | -1.298 | [-2.442, -0.269] | **0.997** | +0.338 | [+0.078, +0.623] | **0.998** | 130 |
| Avg. Dose (LSD μg eq.) | +0.301 | [-0.241, +0.895] | 0.860 | -1.534 | [-2.445, -0.697] | **>.999** | +0.399 | [+0.101, +0.720] | **0.995** | 130 |
| ***VCH Task Behavior*** | | | | | | | | | | |
| 75% Threshold | -0.446 | [-1.092, +0.148] | **0.929** | +0.416 | [-0.382, +1.183] | 0.839 | -0.233 | [-0.439, -0.002] | **0.962** | 113 |
| Hit Rate (75% Contrast Trials) | -0.192 | [-0.583, +0.213] | 0.838 | +0.281 | [-0.447, +0.999] | 0.771 | -0.135 | [-0.349, +0.079] | 0.890 | 113 |
| VCH Rate | +0.328 | [-0.205, +0.918] | 0.887 | -1.589 | [-2.705, -0.537] | **0.999** | +0.504 | [+0.110, +0.946] | **0.996** | 113 |
| ***HGF Estimates*** | | | | | | | | | | |
| Prior Weighting (ν) | +0.229 | [-0.206, +0.649] | 0.859 | -0.667 | [-1.517, +0.094] | **0.946** | +0.259 | [-0.030, +0.588] | **0.964** | 113 |
| Decision Precision (β) | -0.343 | [-0.832, +0.158] | **0.915** | +0.912 | [+0.153, +1.691] | **0.989** | -0.292 | [-0.499, -0.086] | **0.992** | 113 |
| Contingency Belief Evolution Rate (ω) | +0.182 | [-0.331, +0.690] | 0.766 | +0.412 | [-0.376, +1.223] | 0.838 | -0.012 | [-0.297, +0.294] | 0.531 | 113 |

**Supplementary Table S6: Regression table of univariate Bayesian Regression Models.** Statistics refer to regressions comparing beta coefficients (first columns) for predicting (**a**) mean probability of SP-associated PPA history and (**b**) mean number of visual CAPS items endorsed in the past month to the expected change (Δ) in both variables from a 1 SD increase in the predictor (far right columns). For (**b**) only, center columns also summarize the beta coefficient for the hurdle (“hu”) component determining the probability of there being 0 CAPS symptoms—negative values indicate a reduced probability of 0 symptoms. The probability of a coefficient or a Δ being non-zero is the primary value used for statistical inference; it is the equivalent to a p-value. For PPA history variables that do not include a hurdle component, the probability of these two events should be identical—differences are due to probabilistic noise in simulations. HDI = “Highest density interval”—the equivalent of a 94% confidence interval around the mean estimate. See “Bayesian regression model specification and inference approach” in **Supplementary Materials** for more details.

**a**

| **Path** | **β** | **β 94% HDI** | **P(β≠0)** | **Δ** | **Δ 94% HDI** | **P(Δ≠0)** |
| --- | --- | --- | --- | --- | --- | --- |
| ***VCH Decision Noise (β)*** | | | | | | |
| a | -0.031 | [-0.194, +0.146] | 0.639 | -0.016 | [-0.097, +0.073] | 0.639 |
| b | -0.827 | [-1.533, -0.138] | **0.989** | -0.084 | [-0.158, -0.010] | **0.989** |
| c′ | -1.247 | [-2.080, -0.499] | **>.999** | -0.131 | [-0.222, -0.049] | **>.999** |
| NIE | — | — | — | +0.002 | [-0.012, +0.018] | 0.629 |
| NDE | — | — | — | -0.129 | [-0.217, -0.049] | **0.999** |
| Total | — | — | — | -0.126 | [-0.214, -0.045] | **0.999** |
| PMed | — | — | — | -0.015 | [-0.193, +0.113] | 0.628 |
| ***VCH Top-down Bias (ν)*** | | | | | | |
| a | +0.172 | [-0.160, +0.514] | 0.836 | +0.050 | [-0.050, +0.159] | 0.836 |
| b | +0.218 | [-0.456, +0.907] | 0.729 | +0.020 | [-0.042, +0.081] | 0.729 |
| c′ | -1.198 | [-1.984, -0.431] | **0.999** | -0.127 | [-0.214, -0.045] | **0.999** |
| NIE | — | — | — | +0.001 | [-0.008, +0.013] | 0.643 |
| NDE | — | — | — | -0.126 | [-0.212, -0.044] | **0.999** |
| Total | — | — | — | -0.124 | [-0.208, -0.039] | **0.999** |
| PMed | — | — | — | -0.010 | [-0.143, +0.083] | 0.642 |
| ***VCH Response Rate*** | | | | | | |
| a | +0.155 | [-0.131, +0.422] | 0.856 | +0.006 | [-0.005, +0.017] | 0.856 |
| b | +1.224 | [+0.259, +2.288] | **0.995** | +0.098 | [+0.030, +0.163] | **0.995** |
| c′ | -1.336 | [-2.131, -0.531] | **0.999** | -0.140 | [-0.228, -0.054] | **0.999** |
| NIE | — | — | — | +0.006 | [-0.007, +0.021] | 0.838 |
| NDE | — | — | — | -0.134 | [-0.217, -0.052] | **0.999** |
| Total | — | — | — | -0.125 | [-0.214, -0.048] | **0.998** |
| PMed | — | — | — | -0.048 | [-0.243, +0.069] | 0.836 |
| ***VCH 75% Detection Threshold*** | | | | | | |
| a | +0.151 | [-0.012, +0.316] | **0.958** | +0.076 | [-0.006, +0.158] | **0.958** |
| b | -0.894 | [-1.686, -0.151] | **0.987** | -0.093 | [-0.178, -0.011] | **0.987** |
| c′ | -1.102 | [-1.919, -0.345] | **0.997** | -0.116 | [-0.208, -0.033] | **0.997** |
| NIE | — | — | — | -0.011 | [-0.033, +0.003] | **0.941** |
| NDE | — | — | — | -0.113 | [-0.200, -0.032] | **0.997** |
| Total | — | — | — | -0.128 | [-0.216, -0.042] | **>.999** |
| PMed | — | — | — | +0.091 | [-0.040, +0.305] | **0.941** |

**b**

| **Path** | **β** | **β 94% HDI** | **P(β≠0)** | **Δ** | **Δ 94% HDI** | **P(Δ≠0)** |
| --- | --- | --- | --- | --- | --- | --- |
| ***VCH Decision Noise (β)*** | | | | | | |
| a | -0.268 | [-0.466, -0.062] | **0.994** | -0.134 | [-0.233, -0.031] | **0.994** |
| b | -0.283 | [-0.822, +0.250] | 0.861 | -0.193 | [-0.385, +0.014] | **0.958** |
| b (hu) | +0.644 | [-0.120, +1.456] | **0.941** | — | — | — |
| c′ | +0.318 | [-0.251, +0.939] | 0.870 | +0.446 | [+0.078, +0.796] | **0.995** |
| c′ (hu) | -1.603 | [-2.635, -0.613] | **>.999** | — | — | — |
| NIE | — | — | — | +0.058 | [-0.015, +0.166] | **0.957** |
| NDE | — | — | — | +0.436 | [-0.000, +0.760] | **0.991** |
| Total | — | — | — | +0.526 | [-0.000, +0.847] | **0.996** |
| PMed | — | — | — | +0.115 | [-0.034, +0.417] | **0.954** |
| ***VCH Top-down Bias (ν)*** | | | | | | |
| a | +0.197 | [-0.189, +0.584] | 0.836 | +0.054 | [-0.049, +0.175] | 0.836 |
| b | +0.190 | [-0.229, +0.645] | 0.831 | +0.195 | [-0.048, +0.476] | **0.948** |
| b (hu) | -0.648 | [-1.521, +0.214] | **0.926** | — | — | — |
| c′ | +0.356 | [-0.223, +0.955] | 0.900 | +0.498 | [+0.157, +0.866] | **0.998** |
| c′ (hu) | -1.732 | [-2.785, -0.774] | **>.999** | — | — | — |
| NIE | — | — | — | +0.018 | [-0.038, +0.138] | 0.811 |
| NDE | — | — | — | +0.490 | [-0.001, +0.815] | **0.994** |
| Total | — | — | — | +0.529 | [-0.000, +0.875] | **0.996** |
| PMed | — | — | — | +0.038 | [-0.083, +0.308] | 0.811 |
| ***VCH Response Rate*** | | | | | | |
| a | +0.320 | [+0.012, +0.626] | **0.971** | +0.012 | [-0.000, +0.023] | **0.971** |
| b | +0.290 | [-0.233, +0.865] | 0.871 | +0.398 | [+0.066, +0.785] | **0.995** |
| b (hu) | -1.508 | [-2.699, -0.430] | **0.998** | — | — | — |
| c′ | +0.361 | [-0.215, +0.971] | 0.899 | +0.477 | [+0.124, +0.837] | **0.996** |
| c′ (hu) | -1.683 | [-2.778, -0.718] | **>.999** | — | — | — |
| NIE | — | — | — | +0.045 | [-0.012, +0.139] | **0.962** |
| NDE | — | — | — | +0.456 | [-0.001, +0.776] | **0.990** |
| Total | — | — | — | +0.525 | [-0.001, +0.856] | **0.995** |
| PMed | — | — | — | +0.092 | [-0.036, +0.327] | **0.958** |
| ***VCH 75% Detection Threshold*** | | | | | | |
| a | -0.137 | [-0.340, +0.053] | **0.903** | -0.068 | [-0.170, +0.026] | **0.903** |
| b | -0.440 | [-1.134, +0.144] | **0.930** | -0.176 | [-0.361, +0.016] | **0.948** |
| b (hu) | +0.321 | [-0.494, +1.165] | 0.769 | — | — | — |
| c′ | +0.388 | [-0.167, +1.023] | **0.923** | +0.482 | [+0.161, +0.830] | **>.999** |
| c′ (hu) | -1.733 | [-2.745, -0.744] | **>.999** | — | — | — |
| NIE | — | — | — | +0.022 | [-0.021, +0.106] | 0.857 |
| NDE | — | — | — | +0.496 | [-0.000, +0.819] | **0.997** |
| Total | — | — | — | +0.543 | [-0.001, +0.889] | **0.998** |
| PMed | — | — | — | +0.044 | [-0.056, +0.213] | 0.857 |

**Supplementary Table S7: Regression table of multivariate Bayesian mediation models**. For the relationship between (**a**) age of first SP use and history of SP-associated PPAs and (**b**) average SP dose and past-month CAPS visual item endorsements, summary statistics of each path in a standard mediation diagram are listed beneath each potential mediating variable explored. The paths are as follows; a: SP use → mediator, b: mediator → PPA variable, c’: SP use → PPA variable while controlling for the mediator, NIE: the “natural indirect” effect or “mediated” effect of SP use → mediator → PPA variable, NDE: the combined “natural direct effect” of both c’ and (for CAPS vision models only) the hurdle “hu” c’, “Total”: the combined NIE and NDE effects of SP use → PPA variable, “PMed”: the proportion of the Total effect attributable to the NIE. Paths that include “hu” capture the “hurdle” component—how the predictor/mediator shifts the probability of having zero CAPS visual endorsements, such that negative indicates high probability of nonzero symptoms. Δ represents the expected change in a PPA variable from a 1 SD increase in the predictor or mediator. For CAPS vision models, Δ estimates account for both change in mean number of symptoms (β) and the probability of symptoms being nonzero (“hu β” or the hurdle component). For PPA history models without a hurdle component, the probability that β is nonzero is identical to the probability that Δ is nonzero. The probability of a β coefficient or Δ being non-zero is the primary value used for statistical inference; it is the equivalent to a p-value. HDI = “Highest density interval”—the equivalent of a 94% confidence interval around the mean estimate. See “Bayesian regression model specification and inference approach” in **Supplementary Materials** for more details.

**Supplementary Figures**

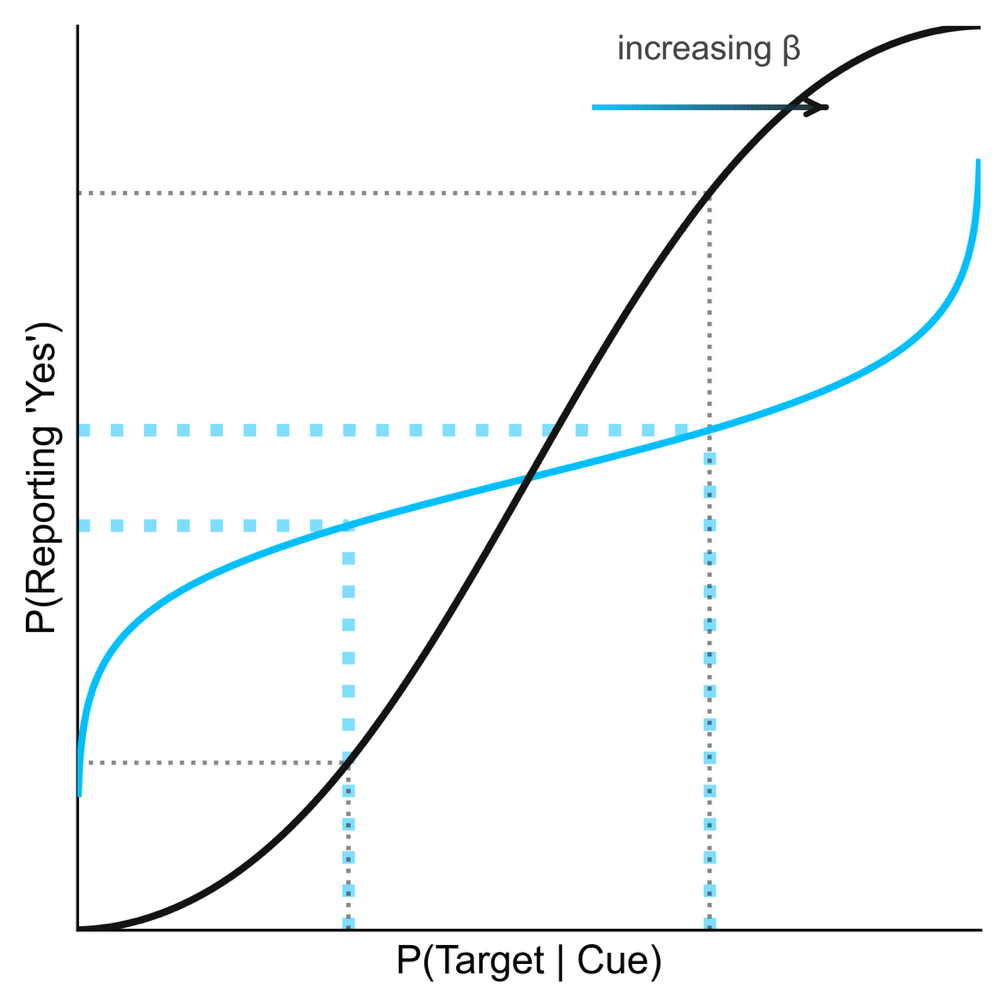

**Supplementary Figure S1**: **Graphical depiction of response probabilities at different decision precision (*β*) values of the Hierarchical Gaussian Filter (HGF).** Both lines depict the unit square sigmoid function (see **Supplementary Equation S1) that** transforms the believed probability of a target being present (*b_t_*) into a response (“yes I see the target” or “no I do not see the target”) probability. The light blue line represents the sigmoid steepness at *β* = 0.5 while the dark gray is *β* = 1.5; lower *β* values bias response probability towards randomness (50%).

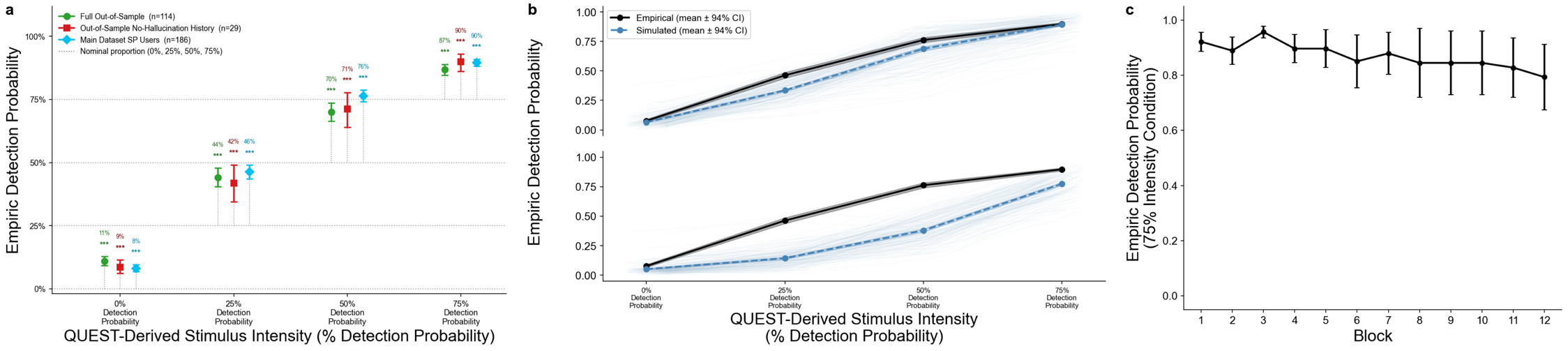

**Supplementary Figure S2**: **Assessment of deviation between QUEST-targeted detection probabilities and empiric detection probabilities in VCH task.** (**a**) Group mean detection probability (responding “Yes, I see it”) over all trials of a given, QUEST-estimated detection threshold. Both green and red points are selected from another online cohort^10^ with a variety of hallucinatory experiences. Green is the full cohort (n=114) while red is a subset of only those denying both auditory and visual hallucinations (n=29)—as assessed using the Chicago Hallucination Assessment Test (CHAT).^20^ The electric blue sample is the current dataset, excluding non-SP users. (**b**) posterior predictive checks for the main dataset (SP users only; n=160; see **Supplementary Materials** “Hierarchical Gaussian Filter (HGF) Model: Bayesian Workflow for Computational Psychiatry”) using empiric (**top**) versus nominal, QUEST-derived (**bottom**) detection probabilities. (**c**) Change (or lack thereof) in detection rate in the 75% contrast condition across all the entire task for the non-hallucinating sample (n=29). All bars represent bootstrapped 94% confidence intervals with 10,000 iterations.

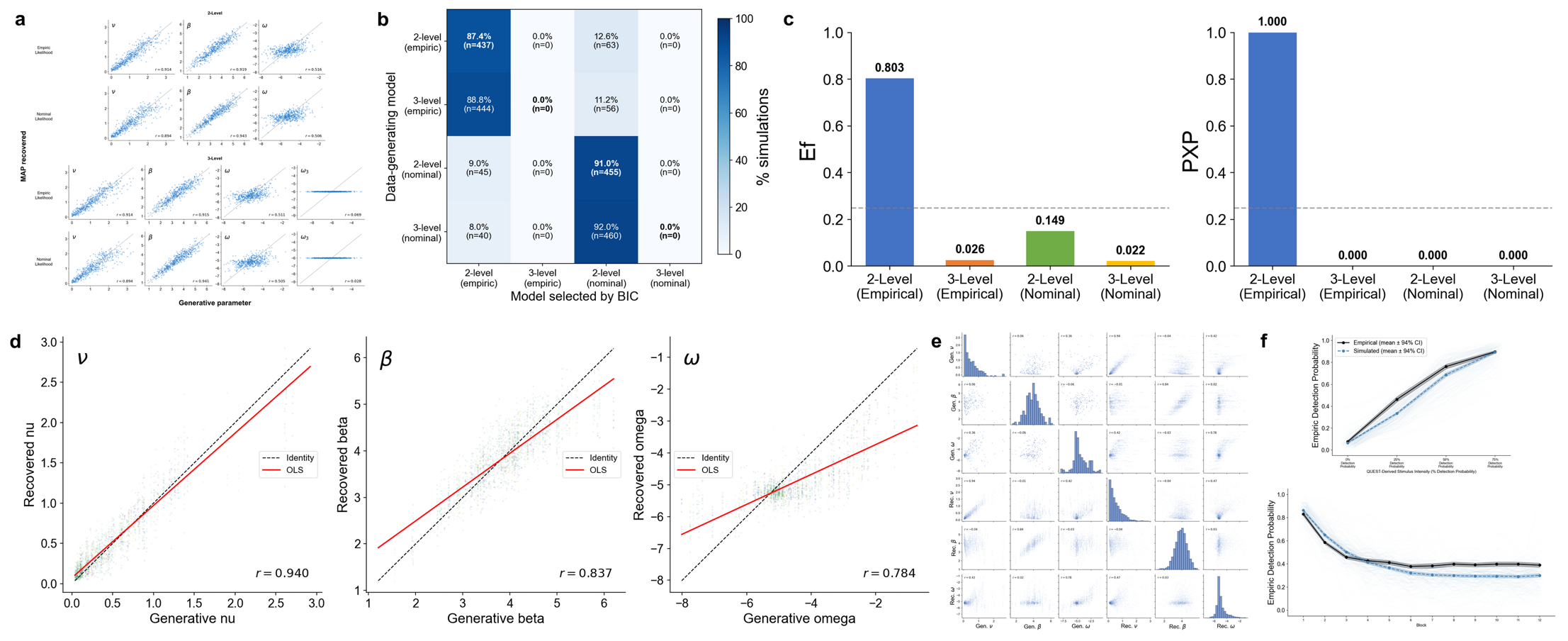

**Supplementary Figure S3: Bayesian workflow for HGF model selection and validation.** See Hierarchical Gaussian Filter (HGF) Model: Bayesian Workflow for Computational Psychiatry” in the **Supplementary Materials** for description of the HGF and models used in comparison. See an excellent example^21^ of Bayesian workflow for further rationale/details of methods that follow. (**top**-**row**) Prior-based parameter recovery, model identifiability, and Bayesian model selection. (**a**) Correlation plots between parameter values sampled randomly from their prior distributions used to generate synthetic data and the median of the recovered parameter value’s posterior distribution, showing excellent recovery for *ν* and *β* (r > 0.9) and acceptable recovery for *ω* (r ~ 0.5), with *ω_3_* demonstrating unrecoverability (r < 0.1). (**b**) A confusion matrix showing the model identifiability via the proportion of generative samples (out of 500) in which each model achieved the lowest BIC following inversion—ideal identifiability would be shaded squares from upper left down, diagonally, indicating that the best-fitting model was the one that had generated the data. Results demonstrate that, regardless of detection probabilities used for likelihoods, the 2-level models were consistently preferred to the 3-level models, most likely indicating the absence of information provided by the third level and the unrecoverable *ω_3_* parameter. (**c**) Random-effects Bayesian model selection (RFX-BMS) results across the four candidate models. The 2-level HGF with empiric detection likelihoods selected based on the greatest expected posterior frequency (Ef = 0.805) and posterior exceedance probability (PXP = 1.00). Bayesian Omnibus Risk (BOR) was 0.00, confirming that Ef varied between models. (**bottom**-**row**) Posterior-based model validation. (**d**) correlation plots between median participant-level parameter estimates used as generative parameters and recovered median estimates following simulation of 10 synthetic behavioral datasets per participant, demonstrating excellent recovery (r = 0.936, 0.832, and 0.764 for ν, β, and ω, respectively), (**e**) a parameter identifiability matrix showing correlations between each generative parameter and all recovered parameters, with the largest off-diagonal correlation observed between generative *ν* and recovered *ω* (r = 0.43) and no correlation between generative *ν* and recovered *β* (r = −0.04) nor generative *β* and recovered *ν* (-0.01), and (**f**) posterior predictive checks (PPCs) showing individual (light, spaghetti lines) and average (dark blue; 94% HDI indicated by shading) model-simulated behavioral trajectories derived from each participants’ median posterior parameter estimates alongside observed data. The model recapitulated increasing detection probability with stimulus strength and decaying detection probability over the course of the experiment. HGF-simulated data slightly underestimated detection probabilities in the 25% and 50% intensity conditions, and estimated slower and greater unlearning of the stimulus-cue association.

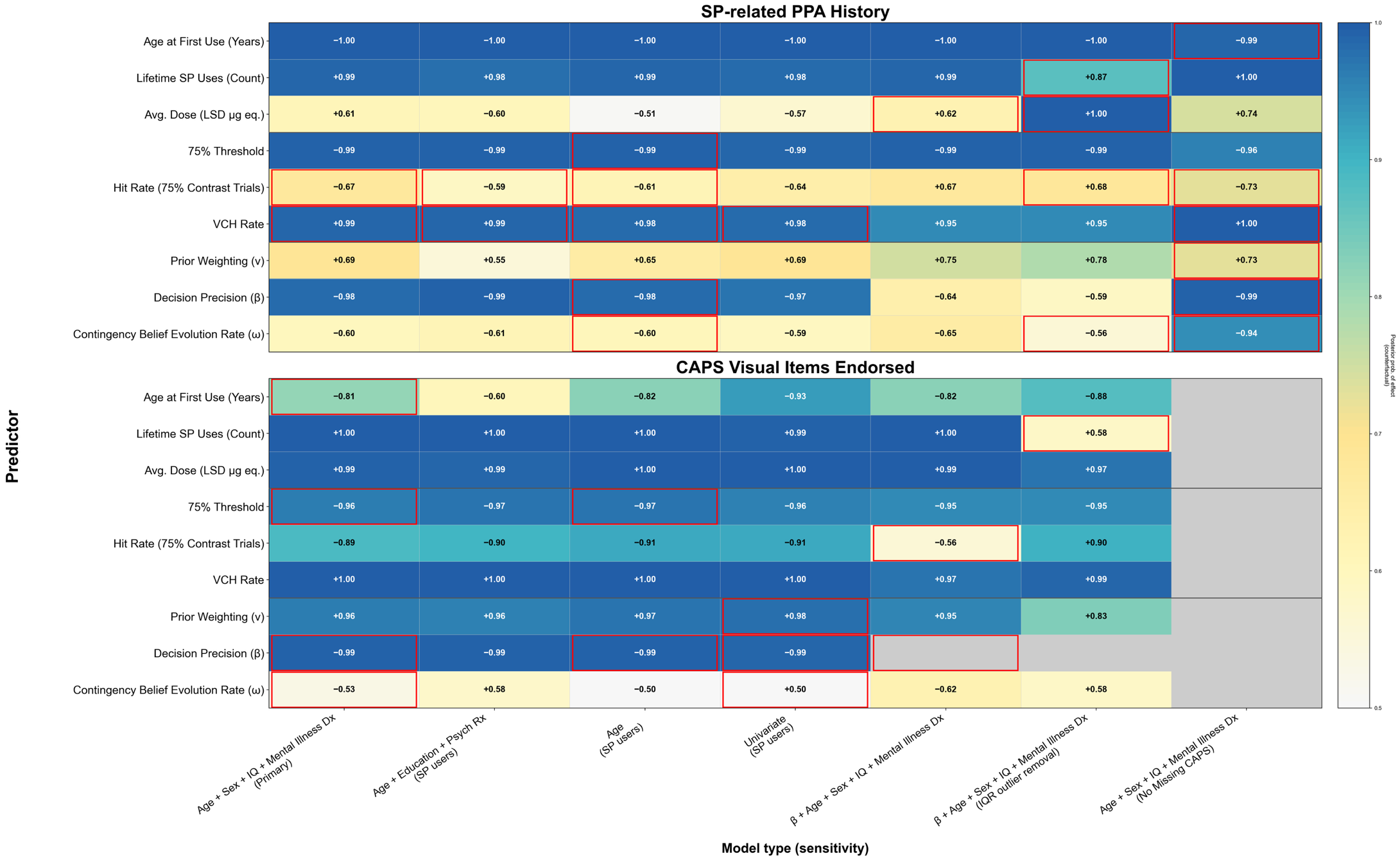

**Supplementary Figure S4: Sensitivity analyses for univariate Bayesian regression models.** For each predictor discussed in the primary manuscript (y-axis), the posterior probability of a 1 SD increase altering (**top**) the probability of having had SP-associated PPAs or (**bottom**) number of visual items endorsed on the CAPS is depicted within the cell for each model type (x-axis). The first column is the model used for figures presented in the manuscript. Subsequent models include: 1) adding 42 SP-naïve participants, 2) removing outlier observations more than 150% of the IQR of the predictor variable, 3) controlling for clinical/demographic variables with some evidence of association with PPA history or CAPS vision items (see **Table 1** and **Supplementary Table 4**), 4) only controlling for age, and 5) no covariates. Cells outlined in red indicate models with one or more significant patterns in scaled residuals from the DHARMa package in R, suggesting possible model misspecification. Grayed cells represent models that were not run.

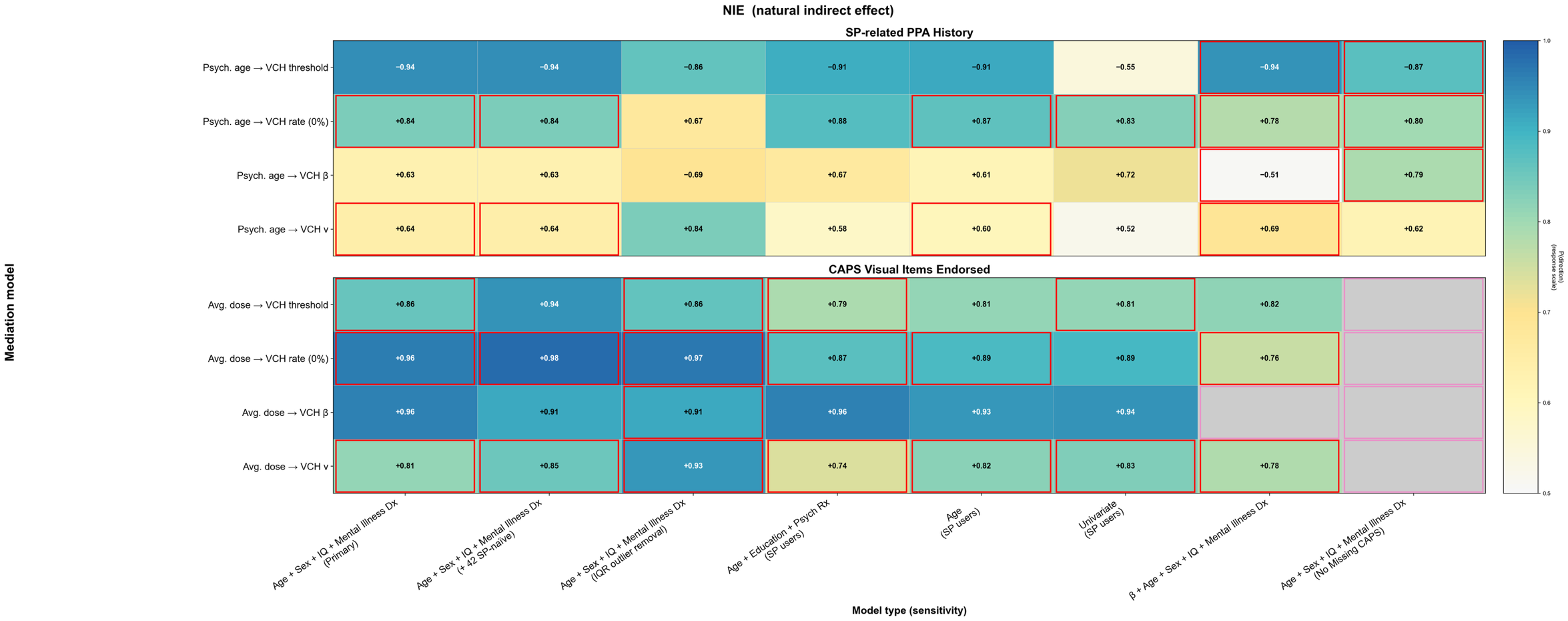

**Supplementary Figure S5**: **Sensitivity analyses for multivariate Bayesian mediation models—mediated path.** For each mediator discussed in the primary manuscript (y-axis), the posterior probability of a 1 SD increase in (**top**) age of first SP use increasing the probability of having had SP-associated PPAs via that mediator or (**bottom**) average SP dose increasing number of visual items endorsed on the CAPS for that mediator is depicted within the cell for each model type (x-axis). The first column is the model used for figures presented in the manuscript. Subsequent models include: 1) adding 42 SP-naïve participants, 2) removing outlier observations more than 150% of the IQR of the predictor variable, 3) controlling for clinical/demographic variables with some evidence of association with PPA history our CAPS vision items, 4) only controlling for age, and 5) no covariates. Cells outlined in red indicate models with one or more significant patterns in scaled residuals from the DHARMa package in R, suggesting possible model misspecification.

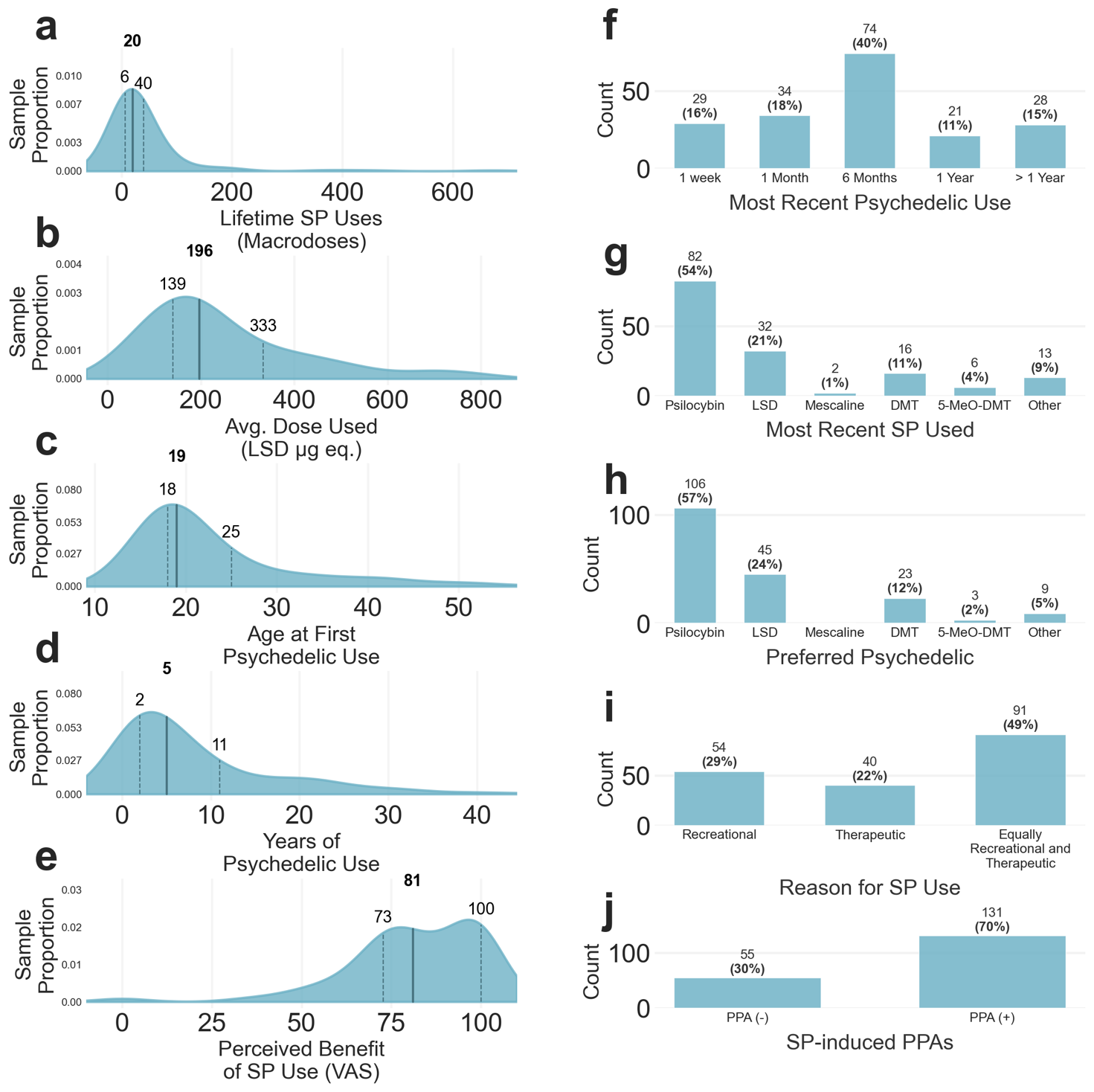

**Supplementary Figure S6.** **SP use history.** Density plots display the distributions of (**a**) number of lifetime SP uses with detectable psychoactive effects (i.e. excluding microdoses), (**b**) average SP dose used in LSD µg equivalents based on subjective intensity, age at first SP use, (**c**) years of SP use (**d**), and (**e**) perceived benefit of SP use measured via visual analog scale (VAS). Dark lines and bolded values indicate medians while dotted lines and non-bold labels indicate 25th and 75th percentile values. (n = 186) Bar charts display the distributions of participants’ (**f**) time since last SP use, (**g**) SP used most recently, (**h**) most used SP, (**i**) primary reason for psychedelic use, and (**j**) history of any PPAs attributed to SP use.

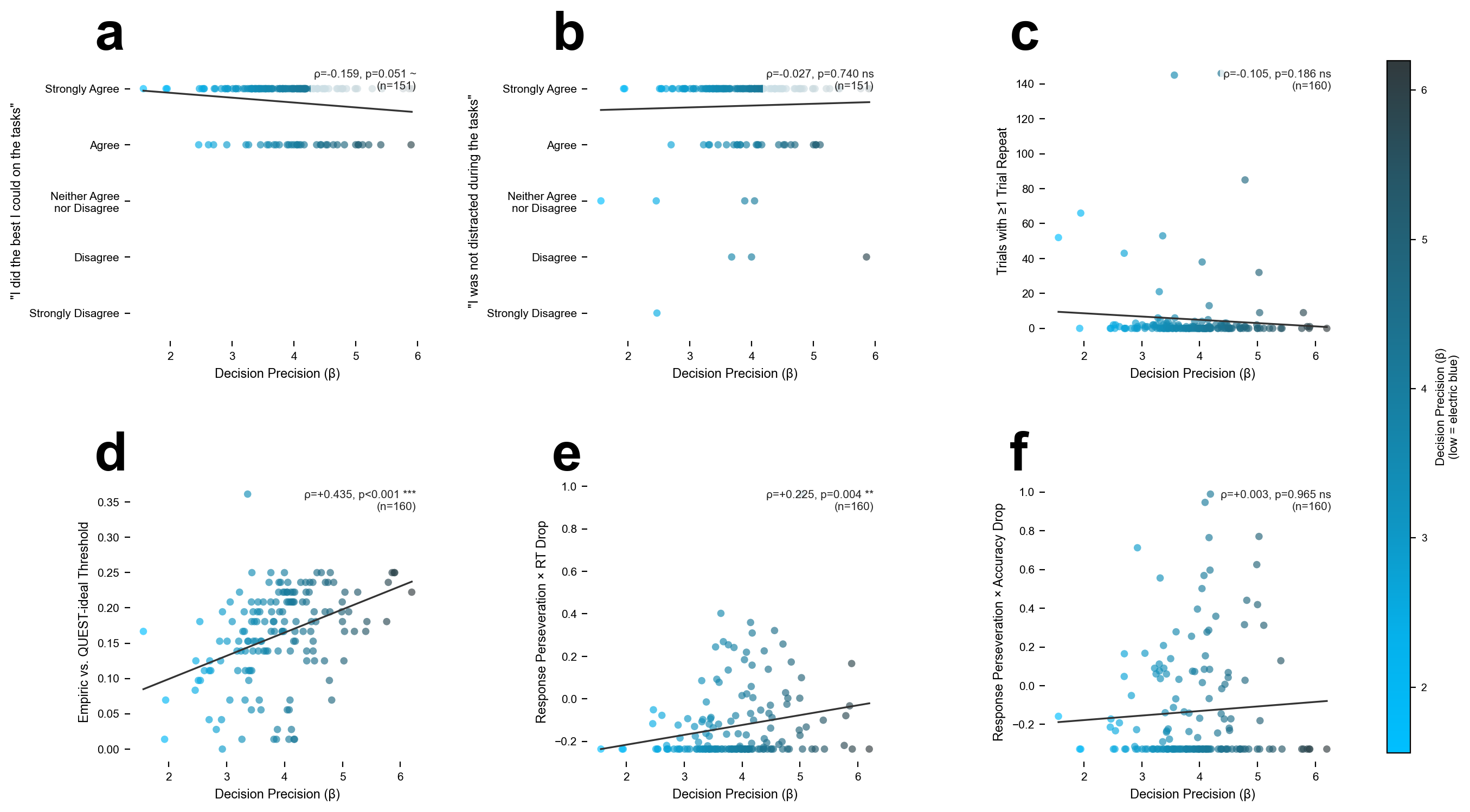

**Supplementary Figure S7**: **Correlations between poor task-engagement indicators and decision precision (*β*).** In order to test whether decreased decision precision reflects poor task engagement (i.e. marks a nuisance confound), we examined the correlation between *β* and (**a**) self-reported effort, (**b**) self-reported distraction during the tasks, (**c**) the number of trials where the stimulus had to be re-presented due to non-response the first time, (**d**) the error in QUEST-estimated visual threshold and empiric probability of detection in the 75% intensity trials, (**e**) a composite score testing for random responses in the form of longest contiguous streak of identical responses (perseveration) multiplied by decrease in median reaction time—Z scored and (**f**) the same composite score but using a change in accuracy (hit rate + correct reject rate / VCH rate + miss rate). We found no correlations between low β and poor task engagement. The only correlations seen were between *higher* *β* values and QUEST-versus-empiric detection probability and reaction-time-multiplied response perseveration composite score.
